# A novel enhancer of *Agouti* contributes to parallel evolution of cryptically colored beach mice

**DOI:** 10.1101/2021.11.05.467454

**Authors:** T. Brock Wooldridge, Andreas F. Kautt, Jean-Marc Lassance, Sade McFadden, Vera S. Domingues, Ricardo Mallarino, Hopi E. Hoekstra

**Author notes:** Nature Ecology & Evolution*, 4 Crinan Street, London N1 9XW, UK.

## Abstract

Identifying the genetic basis of repeatedly evolved traits provides a way to reconstruct their evolutionary history and ultimately investigate the predictability of evolution. Here, we focus on the oldfield mouse (*Peromyscus polionotus*), which occurs in the southeastern United States, where it exhibits considerable coat-color variation. Dorsal coats range from dark brown in mice inhabiting mainland habitat to near white on the white-sand beaches of the southeastern US, where light pelage has evolved independently on Florida’s Gulf and Atlantic coasts as an adaptation to visually hunting predators. To facilitate genomic analyses in this species, we first generated a high-quality, chromosome-level genome assembly of *P. polionotus subgriseus*. Next, in a uniquely variable mainland population that occurs near beach habitat (*P. p. albifrons*), we scored 23 pigment traits and performed targeted resequencing in 168 mice. We find that variation in pigmentation is strongly associated with a ~2 kb region approximately 5 kb upstream of the *Agouti-signaling protein* (ASIP) coding region. Using a reporter-gene assay, we demonstrate that this regulatory region contains an enhancer that drives expression in the dermis of mouse embryos during the establishment of pigment prepatterns. Moreover, extended tracts of homozygosity in this region of *Agouti* indicate that the light allele has experienced recent and strong positive selection. Notably, this same light allele appears fixed in both Gulf and Atlantic coast beach mice, despite these populations being separated by >1,000km. Given the evolutionary history of this species, our results suggest that this newly identified *Agouti* enhancer allele has been maintained in mainland populations as standing genetic variation and from there has spread to, and been selected in, two independent beach mouse lineages, thereby facilitating their rapid and parallel evolution.

---

To gain a complete picture of adaptation, we strive to understand both the molecular mechanisms and the evolutionary processes underlying trait evolution. On one hand, identifying the molecular basis of phenotypic adaptation can provide an opportunity to learn *how* traits vary – in particular, how specific changes in DNA can affect protein function or expression during development to produce the trait of interest. On the other hand, the evolutionary history of a specific allele can provide insights into *when* and *why* traits evolve. Importantly, an allele may be influenced by a combination of neutral and selective forces, which together explain its current distribution and frequency. Thus, the identification of a causal gene, or better, a small gene region or mutation, can serve as a handle with which to probe both the proximate (how) and ultimate (when/why) drivers of trait variation.

Cases of repeated evolution provide a particularly appealing context for understanding the drivers of adaptation. For example, one can ask: did similar phenotypes arise via the same or different molecular changes? While there are empirical examples of selection from new mutations (e.g., Chan et al., 2010; Kowalko et al., 2013; Linnen et al., 2009), it has been suggested that rapid adaptation, in particular within species, may be fueled by selection on the same alleles from pre-existing genetic variation (e.g., Conte et al., 2012; Jones et al., 2018; Oziolor et al., 2019; reviewed in Barrett and Schluter, 2008).

Moreover, it has been argued that changes in *cis*-regulatory elements may be the primary substrate of adaptation (Carroll, 2008; Stern and Orgogozo, 2009; Wittkopp and Kalay, 2011), although many examples of protein-coding changes (e.g., Hoballah et al., 2007; Projecto-Garcia et al., 2013; Zhen et al., 2012; reviewed in Hoekstra and Coyne, 2007) or combinations of both regulatory and coding changes (e.g., Vickrey et al., 2018) have been identified. Nonetheless, when regulatory change has been implicated in repeated evolution, it is still rare that the causal regions, elements, or mutations have been identified (but see Chan et al., 2010; Lewis et al., 2019). This is in part due to the complexity of gene regulatory landscapes and the relative difficulty in testing the effects of a non-coding allele (Pai et al., 2015). By contrast, coding mutations are generally more amenable to identification and functional validation; therefore, when precise mutations have been shown to drive repeated evolution across species, they most commonly correspond to coding mutations (see Martin and Orgogozo, 2013). Thus, it remains difficult to determine the extent to which similar or different mutations contribute to repeated phenotypic evolution and where in the genome they occur.

Variation in pigmentation has long served as a model for the study of adaptation. At the molecular level, the genes and pathways involved in vertebrate pigmentation have been well characterized (Hoekstra 2006). At the phenotypic level, color can vary dramatically in the wild, can be measured straightforwardly, and can have clear links to fitness (e.g., Caro and Mallarino, 2019). One classic and extreme example of rapid color evolution involves the beach mice found in the southeastern United States. There, local populations have independently evolved light coloration on Florida’s Gulf and Atlantic Coasts from a dark-colored mainland ancestor (Steiner et al., 2009). Previous work identified three genomic regions involved in differences between beach and mainland mice (Steiner et al., 2007), which were subsequently localized to three pigmentation genes: the *Melanocortin-1 receptor* (*Mc1r*), *Agouti signaling protein* (*ASIP*), and *Corin* (Hoekstra et al., 2006; Manceau et al., 2011; Manceau et al., in review). In mammals, the interaction between *Mc1r* and its inverse agonist *Agouti* mediates the switch from dark (eumelanin) to light pigment (pheomelanin) production (Barsh, 1996; Ollmann et al., 1998; Vrieling et al., 1994), while their interaction with *Corin*, in turn, can mediate the fine-tuning of pigment patterns (Enshell-seijffers et al., 2008; Manceau et al., in review). In Gulf Coast beach mice, a single missense mutation in *Mc1r*, together with *cis*-regulatory changes in *Agouti* (as well as *Corin*) together are largely responsible for their light pigmentation. Therefore, changes in genes at multiple levels of the pigment pathway have been implicated in the evolution of camouflaging coloration in Gulf Coast beach mice.

In contrast, the genes (and mutations) contributing to light coats of the Atlantic coast beach mice have remained elusive. For example, the *Mc1r* amino acid change found in Gulf Coast mice is absent from Atlantic coast mice (Hoekstra et al., 2006), and no new *Mc1r* mutations are associated with color variation or have a measurable effect on Mc1r function (Steiner et al., 2009). In addition, there are no differences in the *Agouti* coding region between mainland mice and beach mice on either coast (Steiner et al., 2007). While this implies a likely role for regulatory changes, the precise regulatory elements(s) and mutation(s) driving *cis*-acting differences in *Agouti* expression in Gulf beach mice, and possibly Atlantic mice, have not yet been identified. As a result, it remains an outstanding question whether the similarly light-colored Gulf and Atlantic Coast beach mice carry the same or distinct pigmentation alleles.

Here we return to the classic case of adaptation in Gulf and Atlantic Coast beach mice, first described over a century ago (Howell, 1920; Sumner, 1926) and capitalize on naturally occurring color variation in a single mainland population to identify the molecular basis of adaptation, using the first chromosome-level genome assembly for *P. polionotus*. Specifically, we employ an association-mapping approach to identify a ~2 kb previously uncharacterized non-coding region of *Agouti* associated with color variation. We then show that this 2 kb region can drive dermal expression in *Mus* embryos, demonstrating its regulatory activity in the skin during the establishment of pigmentation. Finally, we reveal the evolutionary history of this regulatory element to show both strong selection on the light *Agouti* allele in a phenotypically variable population, and that this same allele is fixed in beach mice of both the Gulf and Atlantic Coasts. Together we find both the molecular basis and evolutionary history differs markedly between two key genes involved in beach mouse adaptation – *Agouti* and *Mc1r* – highlighting that there can be multiple genetic solutions to the same ecological challenge, even within species.

## Results

### Assembly of a high quality, chromosome-level genome for *P. polionotus*

We generated whole-genome sequencing (WGS) data and assembled a *de novo* high-quality reference genome for the oldfield mouse, *Peromyscus polionotus subgriseus* (BioProject number PRJNA494229). The final genome was 2.645 Gb in length with a N50 scaffold length of 13 Mb. We could anchor 97% of the *de novo* assembled bases into 23 autosomes and the X chromosome using high-density genetic linkage maps for *Peromyscus*. Our estimates indicate that the assembly contains 95.4% and 94.8% of single-copy core mammalian and euarchontoglire genes, respectively. Our annotation strategy, which combined comparative *in silico* and evidence-based approaches, identified 18,502 protein-coding genes having orthologs in the *Mus* genome, 536 paralogs of *Mus* genes and 1,912 additional genes showing homology with known proteins from curated databases. This complete, high-quality genome enables evolutionary analyses of genome-wide variation across populations of this species.

### Recent and independent evolution of beach mice on the Gulf and Atlantic coasts

To better estimate the timing and pattern of divergence in the beach and mainland subspecies (Fig. 1A), we sampled six beach and five mainland populations, representing nine of the fourteen extant *P. polionotus* subspecies (Fig. 1B) as well as the closely related sister species, *P. maniculatus nubiterrae*. Using 1000 randomly distributed genome-wide SNPs derived from putatively neutral regions in a targeted sequence-capture dataset, we generated a highly supported phylogeny confirming the independent origin of beach mice in the Gulf and Atlantic coasts from an ancestral mainland form (Fig. 1C), consistent with previous studies (Domingues et al., 2012; Kalkvik et al., 2018; Mullen et al., 2009; Steiner et al., 2009). The Gulf Coast beach mice form a paraphyletic group with adjacent mainland populations, all of which share a common ancestor between 3.5-7.2 thousand years ago (kya). Similarly, the Atlantic Coast beach mice share a common ancestor with their closest mainland counterparts 2.9-6.4 kya, suggesting that both Gulf and Atlantic beach lineages originated at approximately the same time. In general, we find that the relationships of subspecies in the phylogeny mirrors their geographic distribution, a pattern that is supported by a genetic Principal Components Analysis based on genotype likelihoods from all loci in the sequence-capture dataset (gPCA; Fig. 1D). The evolutionary history of both Gulf and Atlantic beach mice as well as several mainland populations provides a demographic context in which to understand the evolution of crypsis.

**Figure 1.**
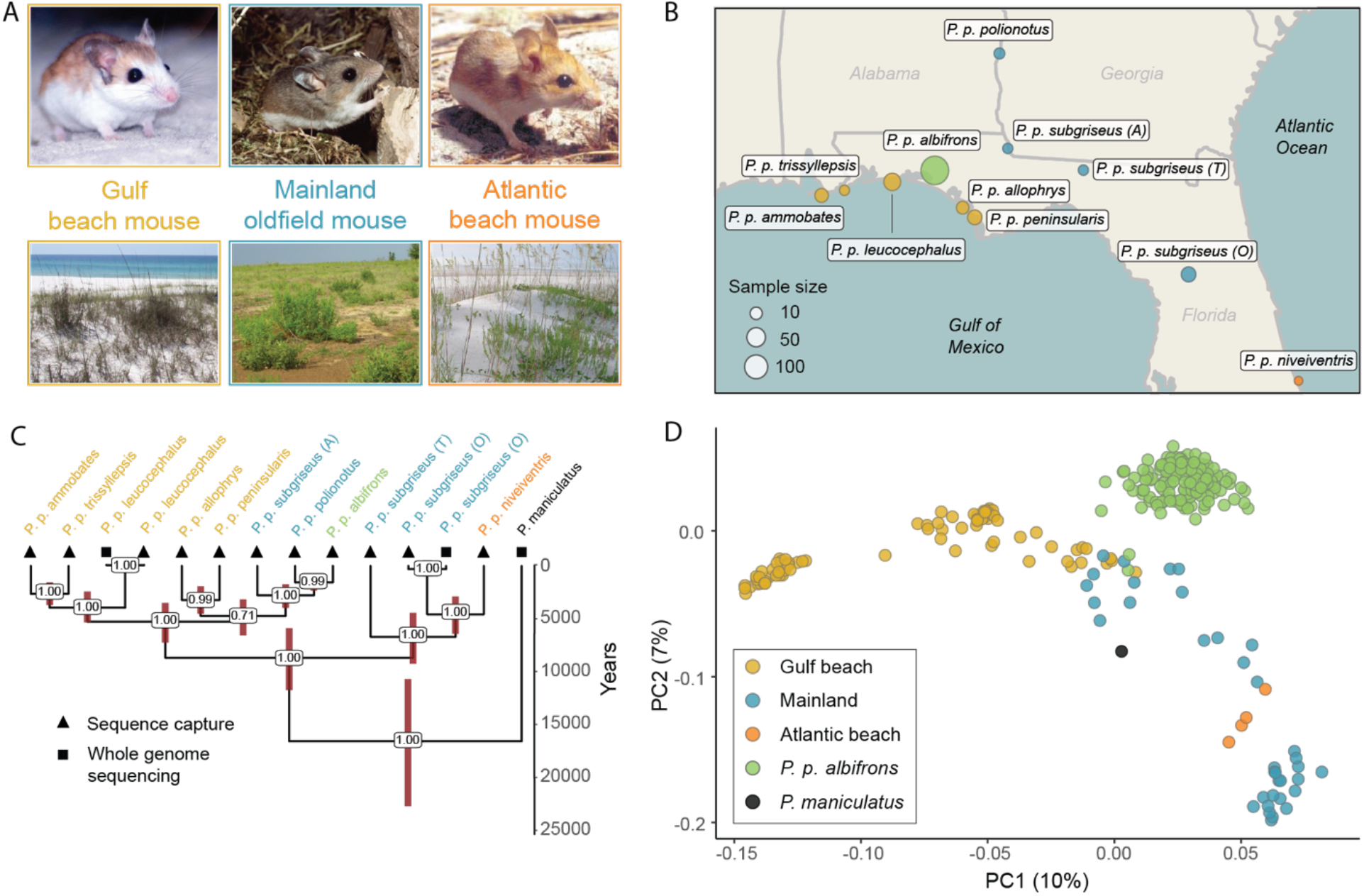
Distribution and relationships of beach and mainland subspecies of *P. polionotus*. **A.** Representative images of beach and mainland subspecies of *P. polionotus* as well as corresponding habitats. **B.** Map of the southeastern United States showing sampling locations of populations included in this study (see Table S1 for details). Sample sizes are indicated in parentheses; the area of each circle corresponds to sample size. *P. p. subgriseus* was sampled from three locations: O = Ocala, A = Apalachee, T = Tall Timbers Research Station. **C.** Time-scaled phylogeny of sampled populations. Numbers at nodes represent bootstrap support; red bars 95% confidence intervals of divergence time. Populations are annotated with one of two sequencing strategies used in this study. **D.** The first two dimensions of a principal component analysis (PCA) based on genotype probabilities; each dot represents an individual with sample sizes given in B.

### Phenotypic variation in a single mainland population (*P. p. albifrons*)

We sampled one mainland population neighboring beach habitat, *P. p. albifrons*, that exhibited a wide range of coat colors – from light and sparsely pigmented coats similar to those of beach mice to the dark and extensively pigmented coats typical of mainland mice. To better characterize and quantify this variation, we measured 23 coat-color traits in 168 skin specimens of *P. p. albifrons* (Fig. 2A). All traits were related to either the distribution of pigmentation (e.g., tail-stripe length) or intensity of pigment (e.g., dorsal hue, brightness) and are known to vary among beach mouse populations (Steiner et al., 2009). To establish reference points with which to compare the *albifrons* population, we scored the same 23 traits in representative mice from Gulf, Atlantic, and mainland subspecies (Table S1).

**Figure 2.**
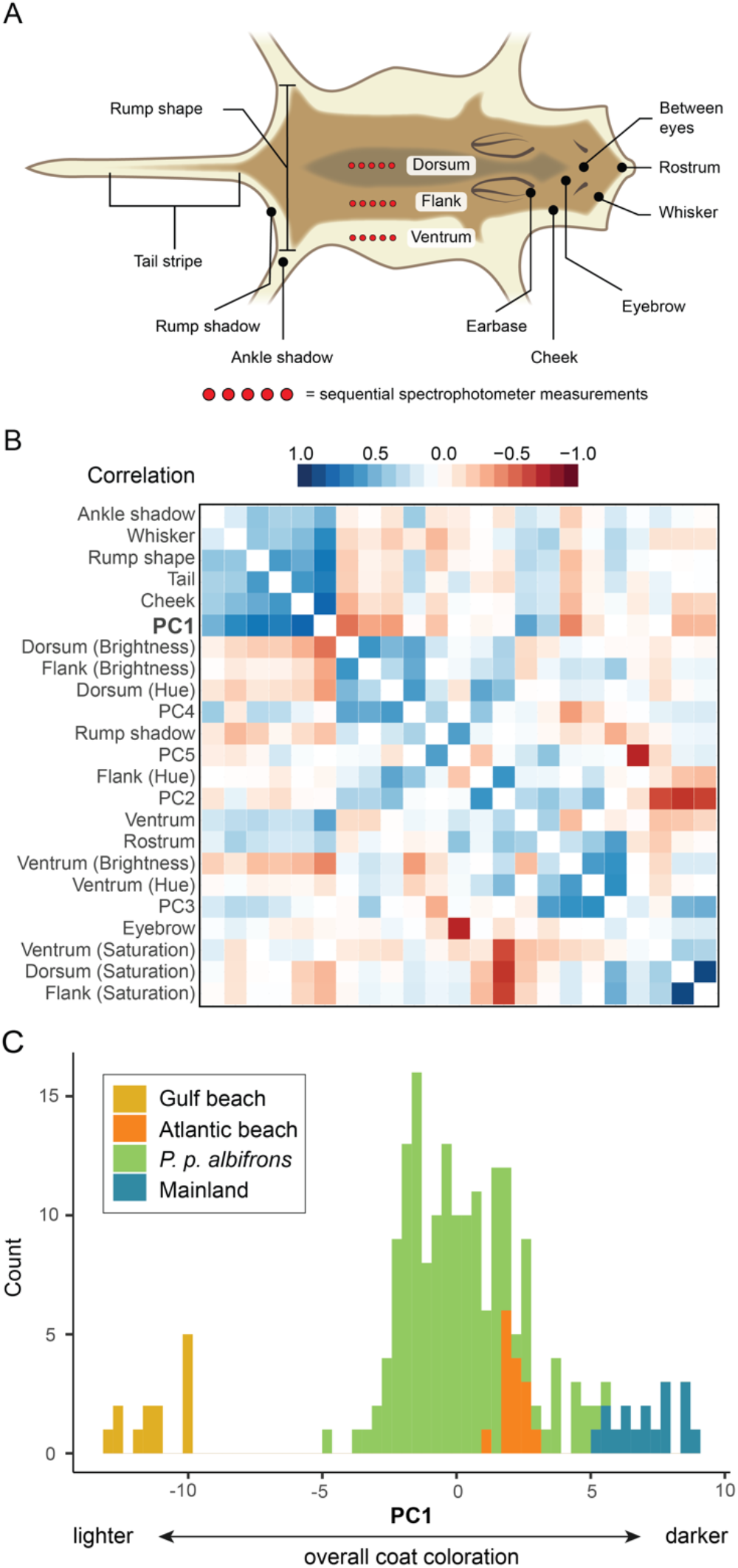
Phenotypic variation in the *P. p. albifrons* population. **A.** Cartoon showing traits used to characterize pigment and pattern variation in museum specimens, categorical scores (black) and spectrophotometric measures (red; see Methods for details). **B.** Pairwise correlation among pigmentation traits and the first 5 phenotypic PCs in *albifrons* mice (n = 168). Color indicates direction and strength of the correlation. Asterisks denote significance: * = p < 0.05, ** = p < 0.01, *** = p < 0.001. PC1 is highlighted by dashed lines. Invariant traits are not shown. **C.** Frequency distribution of PC1 scores for *albifrons* mice as well as representative Gulf (n = 13), Mainland (n = 17), and Atlantic (n = 15) mice.

We found that many pigment traits are highly correlated in the *albifrons* population (Fig. 2B). A principal components analysis (PCA) shows that six traits – dorsal brightness, tail-stripe length and the extent of pigmentation on the cheek, rump, whisker, and ankle – heavily load on phenotypic PC1 (pPC1), and that a specimen’s pPC1 score is a strong predictor of overall lightness or darkness (Fig. 2B, Fig. S1). Remaining traits also form distinct clusters, but none of these additional clusters encompass as many traits as pPC1 or show the same strength of association with overall coloration (Fig. 2B). The highest pPC1 values observed in the population represent the darkest mice, which are similar in appearance (and pPC1 score) to the mainland subspecies, *P. p. polionotus* (Fig. 2C). And while the lightest *albifrons* individuals are still darker than the geographically proximate beach subspecies *P. p. leucocephalus* – the palest form of the Gulf beach mice – many individuals with intermediate pPC1 scores are comparable to a typical Atlantic beach mouse (e.g., *P. p. niveiventris*; Fig. 2C). Despite this range in coloration that encompasses both beach and mainland phenotypes, none of these pigment traits show a significant association with population structure, not surprising given that the *P. p albifrons* population has little detectable genetic structure (Fig. S2).

### Association between pigmentation and a non-coding region of *Agouti*

Capitalizing on the extensive color variation observed within the panmictic *albifrons* population, we performed single-variant association mapping using the sequence-capture data from this population. These data include 6547 putatively neutral biallelic SNPs from across the genome as well as the genomic regions encompassing two pigmentation genes, *Agouti* and *Mc1r* (190kb and 150kb in length, respectively, which includes all exons and known regulatory regions). In our scan, we detected a single region associated with pPC1 that exceeded the genome-wide significance threshold (p<1.23×10^−5^ corrected for number of effective tests; Fig. 3A) in the *Agouti* locus. A closer investigation of this region revealed three SNPs significantly associated with pigment variation, spanning 1,756 bp, in strong linkage disequilibrium (mean r^2^ = 0.85). A single SNP at position chr4:9,845,301 showed a markedly stronger association with pPC1 than the other two (Fig. 3B). This SNP is located between two untranslated exons (exons 1D and 1E), is 120bp upstream of a cluster of SINE elements in reverse orientation relative to the transcription of *Agouti,* and is 5,641 bp upstream of the first coding exon (exon 2). Genotype-phenotype regressions show an additive effect of this locus, which explains 36% of the variance in pPC1, as well as a substantial degree of additive variation in pPC1-correlated traits such as dorsal brightness (19%) or tail stripe length (7.2%) (Fig. 3C). Together, these data point to a small non-coding region of *Agouti,* containing a mutation(s) having a major effect on variation in overall pigmentation in *P. p. albifrons*.

**Figure 3.**
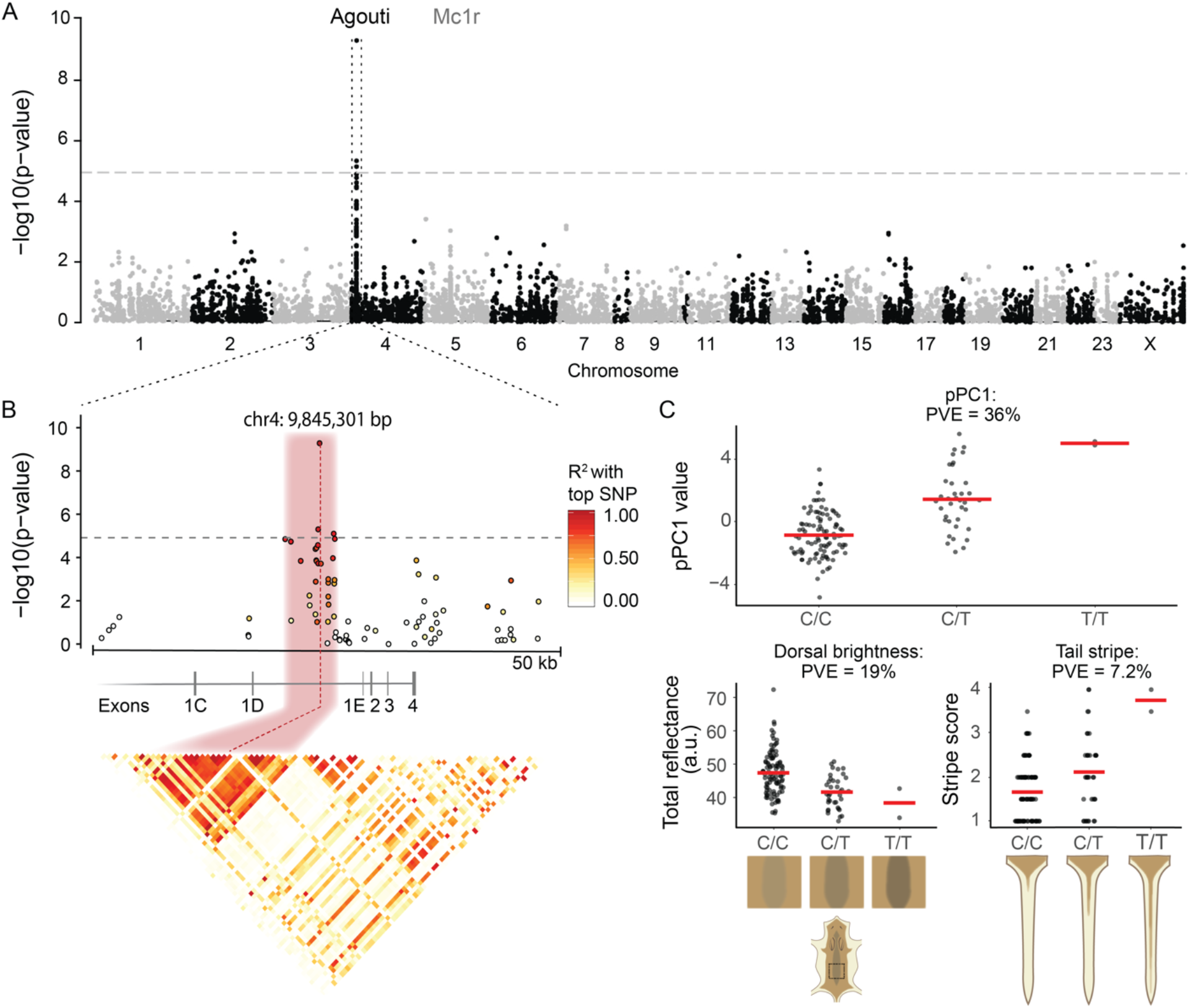
Genotype-phenotype associations in the *Agouti* locus. **A.** A single peak on chromosome 4 that exceeds the genome-wide significance threshold (dashed line) associates with variation in phenotypic PC1. No association observed in *Mc1r*. **B.** Zoom in of the *Agouti* association peak. Dots represent variants; color represents the strength of linkage disequilibrium (LD) with the most highly associated SNP variant at chr4:9,845,301 bp. A single ~2 kb region (pink) shows high levels of LD. Correlation matrix below displays pairwise LD between all variants in the 50 kb region. **C.** Distribution of pPC1 scores (above) and two representative traits (below) by genotype at chr4:9,845,301 bp. Red lines indicate mean trait value by genotype. PVE = percent variance explained. Cartoons illustrate differences in traits by genotype.

### The candidate *Agouti* region is capable of regulatory activity

To determine if this ~2 kb *Agouti* region associated with pPC1 is capable of regulatory activity, we first determined whether the region overlaps with known regulatory elements (Fig. 4A). In the homologous region and ±10 kb surrounding sequence in *Mus,* we observe few known regulatory elements, none of which are associated with dermal tissues (Table S2). Moreover, it does not overlap with any previously identified regions associated with pigment variation in other *Peromyscus* species (e.g., Linnen et al., 2013; Fig. S3). As sequence conservation can be indicative of conserved molecular function, we next examined sequence similarity across 27 rodents in the 5 kb upstream and downstream of the top associated SNP (which also includes the two linked variants and SINE elements). Surprisingly, conservation within rodents was minimal, with only a subset of the species – the superfamily Muroidea – showing greater than 50% sequence similarity for the majority of the region (Fig. 4B; Figure S4). These data suggest that if this region has regulatory function, it is likely to have evolved recently.

**Figure 4.**
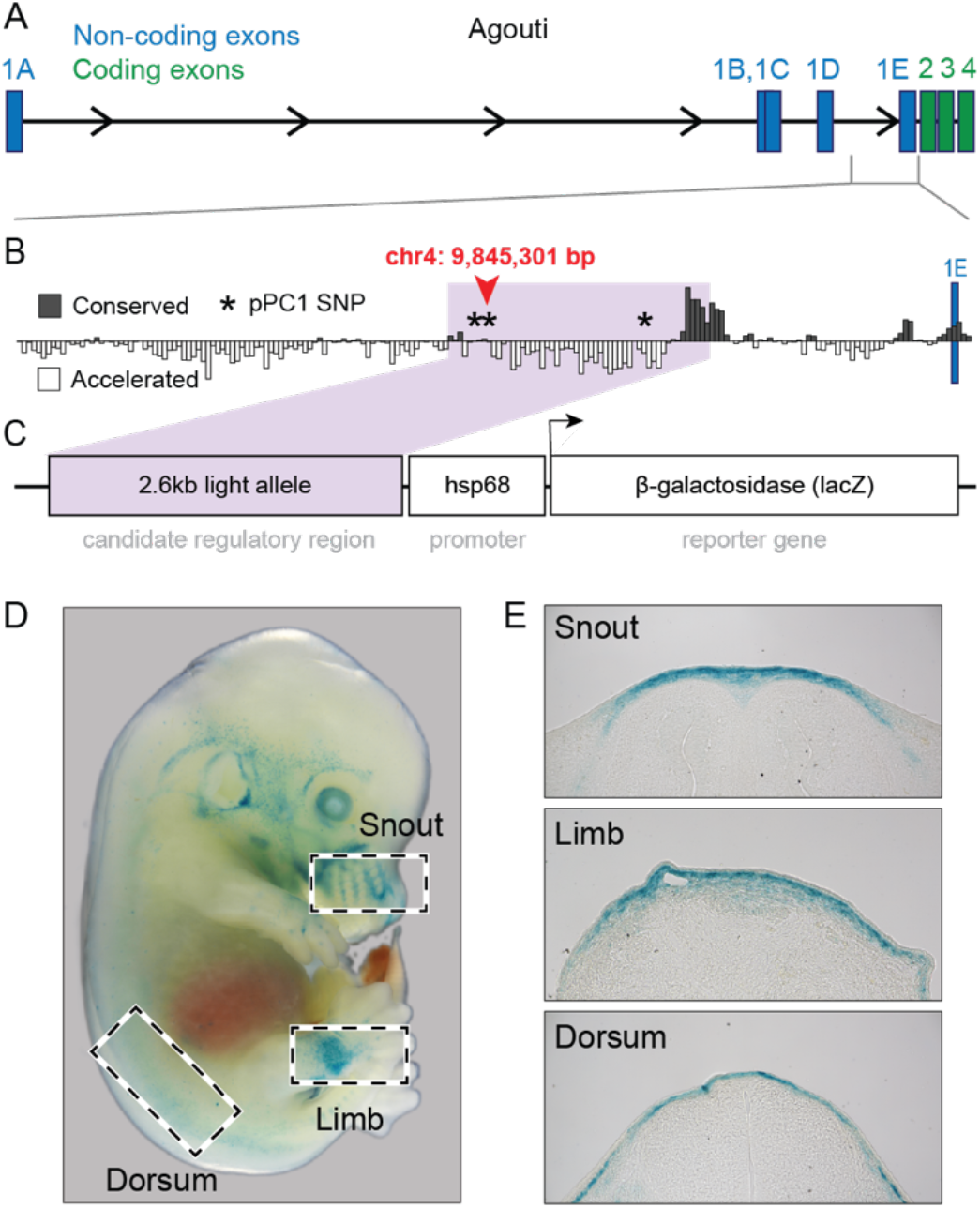
Location, conservation, and activity of candidate *Agouti* regulatory region. **A.** Coding structure of the ~180 kb *Peromyscus Agouti* locus, including non-coding (blue) and coding (green) exons. **B.**PhyloP sequence conservation amongst 27 rodent species in the 10 kb encompassing the SNP most highly associated with pPC1 (chr4:9,845,301); this SNP and 2 others with a significant pPC1 association are denoted by asterisks (*). Purple shading highlights the 2.6 kb region cloned into the *lacZ* reporter plasmid. **C.** Structure of *lacZ* reporter construct. **D.** Stage E14.5 transgenic *Mus* (FVB/NJ) embryo stained for *lacZ* expression (blue). **E.** Three tissue sections showing *lacZ* expression localized to the dermis.

To assess whether the candidate region of *Agouti* contains functional enhancers, we cloned a 2.6 kb sequence that spans 0.5 kb upstream and 2.1 kb downstream of the most strongly associated variant (i.e., chr4:9,845,301) and includes the two additional associated variants (chr4:9,845,152, chr4: 9,846,908) as well as a small downstream region conserved in rodents (Fig. 4B, Fig. S5). We then inserted this sequence upstream of a minimal promoter and *lacZ* reporter gene (Fig. 4C). Given the currently limited transgenic techniques available for *Peromyscus*, the resulting construct was injected into embryos of *Mus* (strain FVB/NJ), and embryos were collected at stage E14.5, a timepoint when *Agouti* expression plays a key role in the establishment of pigment prepatterns in both *Mus* and *Peromyscus* (Manceau et al., 2011). Of the 14 embryos with independent genomic integrations of the *lacZ* construct (verified by PCR), we observed consistent *lacZ* expression in the skin of eight embryos, although expression was spatially variable across embryos (Fig. 4D; Fig. S5). Histological analysis showed that *lacZ* was localized to the dermis, corresponding to the known site of endogenous *Agouti* expression during embryonic development (Fig. 4E). Together, the results of these experiments suggest that this previously undescribed ~2 kb non-coding region contains a *cis*-regulatory element (or possibly multiple elements constituting a *cis*-regulatory module) capable of driving *Agouti* dermal expression during embryonic development.

### The light *Agouti* allele shows a signature of positive selection

We next tested if there was evidence of natural selection acting on the light-associated allele at this regulatory element, which is found at 86% frequency in the *albifrons* population (Table S3). In the region surrounding the top associated SNP (chr4:9,845,301), we found that haplotype homozygosity decays more quickly for the dark allele than for the light allele, a signal consistent with recent positive selection for light pigmentation (Fig. 5A). This signal of elevated homozygosity is statistically significant, with all three candidate SNPs identified in our association analysis (as well as 15 additional SNPs in this region) showing a significantly positive integrated haplotype score (IHS, p < 0.05; Fig. 5B). We did not detect a signal of selection at these candidate SNPs in any other population, although low sample sizes and lack of polymorphisms limit our power. Together, these data support the hypothesis that natural selection, most likely on light coloration, has led to an increase in light *Agouti* allele frequency in the *P. p. albifrons* population.

**Figure 5.**
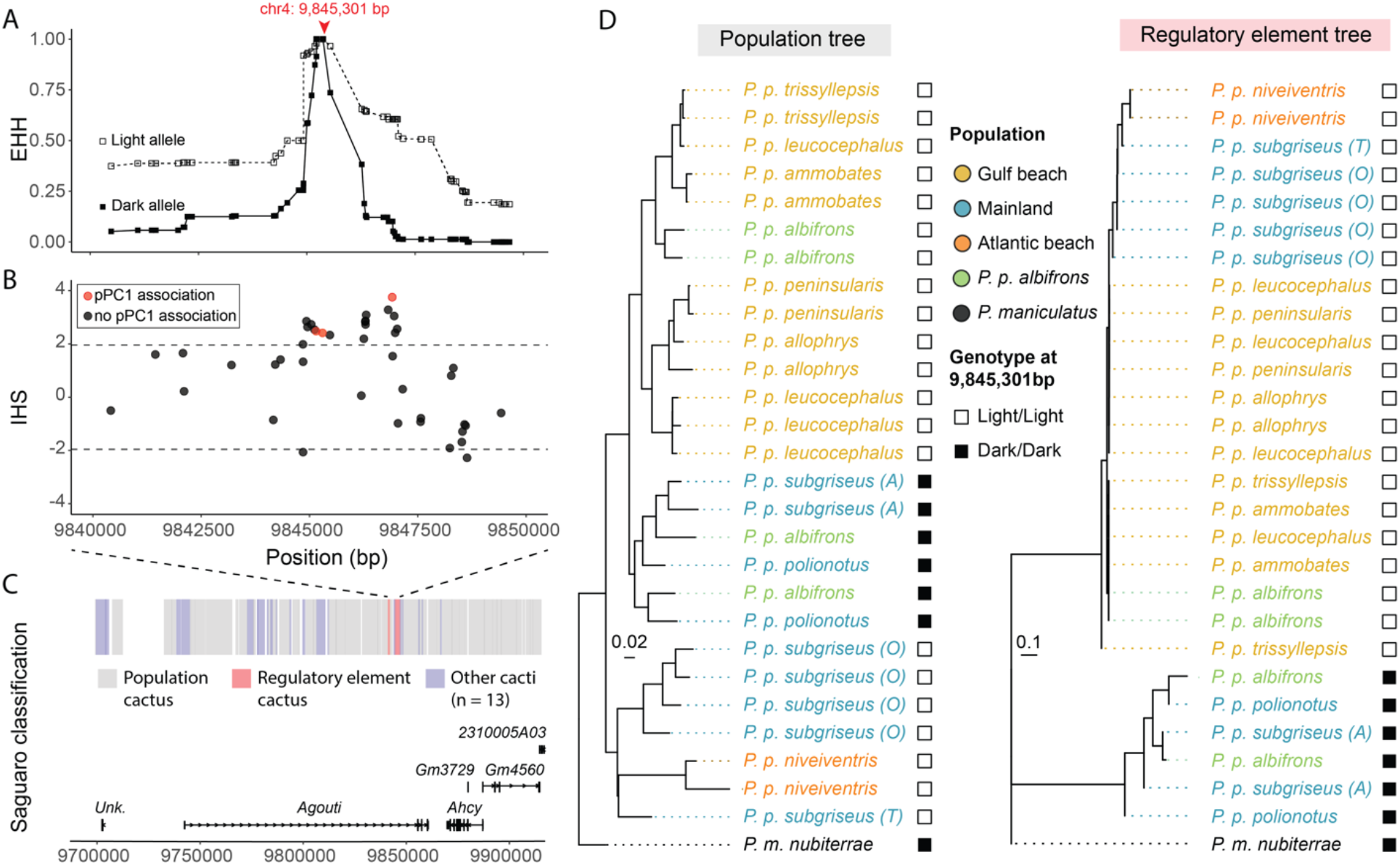
Evolutionary history of the derived light *Agouti* allele. **A.** Extended haplotype homozygosity (EHH) decay plot of 10 kb within the *Agouti* locus, showing that the light allele maintains higher levels of homozygosity than the dark allele around the top-associated SNP (chr4:9,845,301 bp), consistent with a signal of positive selection. **B.** Integrated haplotype score (IHS) calculated for the same *Agouti* region. Values >1.96 indicate statistically significant EHH for the reference allele (p < 0.05). **C.** Saguaro-based classification of local relationships across *Agouti*. Regions that fit to a common single topology (“Population cactus”) are shown in grey. Saguaro independently identified a unique topology (“Regulatory element cactus”) shown in pink, spanning two neighboring regions of 636 bp (position 9,841,443 to 9,842,079 bp) and 2171 bp (9,844,852 to 9,847,023 bp), including the top-associated SNP (chr4:9,845,301 bp). **D.** Cactus topologies using *P. maniculatus* as outgroup. The population cactus topology closely matches the population tree (shown in Fig. 1C), while the regulatory element cactus topology separates individuals homozygous for the light or dark *Agouti* haplotypes, with internal branch lengths suggesting a recent origin of the light *Agouti* allele.

### The light *Agouti* allele is fixed in both Gulf and Atlantic beach mouse populations

Given the evidence for non-neutral evolution at this newly identified *Agouti* regulatory element, we next aimed to infer whether it exhibits a unique evolutionary history relative to the rest of the genome. Using *Saguaro* (Zamani et al. 2013), we generated local phylogenies in variably sized genomic windows across the *Agouti* locus for the combined dataset of beach and mainland populations, and then compared these phylogenies to the population tree constructed from genome-wide neutral loci. At *Agouti*, we find that much of the locus fits a topology (cactus 6) that coarsely mirrors the population-level phylogeny (Fig. 5). In contrast, one unique topology (cactus 4) is exclusively derived from two small regions that include the top-associated SNP and closely match the regulatory element identified in our mapping experiment (i.e., chr4:9,841,443 - 9,842,079 and chr4:9,844,852 - 9,847,023; Fig 5C). This unique topology clusters all individuals homozygous for *Agouti* light alleles into a single clade with short branch lengths, consistent with a recent origin for this allele, while all dark allele individuals fall into a clade with longer branch lengths similar to that observed in the population tree (Fig. 5D). Therefore, not only does this unsupervised approach identify the same region of *Agouti* as was localized in both the independent association and selection analyses, but it also points to a single origin for the light *Agouti* allele. These results show that the light *Agouti* allele arose once and is now shared by both the Gulf and Atlantic beach mice, even though these lineages are geographically distant and independently colonized their respective beach environments from mainland ancestors. This scenario, in which the same allele is inherited from a common ancestor and repeatedly selected for in multiple lineages, suggests that parallel genotypic evolution (sensu Rosenblum et al., 2014) has been an important factor in the evolution of beach mouse coloration.

## Discussion

The question of how adaptation proceeds at the molecular level, and how predictable the process is, has long been of interest to evolutionary biologists. Debate around, for example, the locus of adaptation (i.e., coding versus regulatory mutations), the source of adaptive mutations (i.e., *de novo* mutations versus pre-existing variation) and the repeatability of this process (i.e., do the same or different mutations lead to similar, independently evolved traits) has been lively (e.g., Barrett and Schluter, 2008; Bolnik et al., 2018; Hoekstra and Coyne, 2007; Manceau et al., 2010; Martin and Orgogozo, 2013; Rosenblum et al., 2014; Stern and Orgogozo, 2008). Here, we provide insight into these questions from a classic system, first described over a century ago (Howell 1920): the recent adaptation of two independent lineages of beach mice to novel white-sand habitat through the evolution of camouflaging color. While the *Agouti* gene has been shown to contribute to the evolution of Gulf beach mouse color pattern through changes affecting both its expression level (Steiner et al., 2007) and spatial domain (Manceau et al., 2011), the molecular basis of these regulatory changes remained unclear. Furthermore, without information about the underlying regulatory region(s), the question of whether the genetic basis of light pigment in the Gulf and Atlantic Coast beach mice was the same or different was unknowable. Here, we uncovered a novel regulatory element in the *Agouti* gene and provide evidence that an allele of this element is associated with lighter pigmentation, and that it has been selected repeatedly from standing genetic variation in both lineages of beach mice.

To identify mutation(s) that contribute to changes in pigmentation, we first identified a population (*P. p. albifrons*) that was phenotypically variable, ranging from light beach to dark mainland coat color. While most populations show little variation in pigmentation, this mainland population appears unique, likely because of its geographic proximity (~25 km) to the beach habitat, its patchwork of light sandy and dark loamy soil, and intermediate level of vegetative cover relative to the open beach and dense mainland oldfield habitats. By conducting genetic association mapping in this variable population, we were able to narrow in on a small – approximately 2 kb – non-coding region which is strongly associated with overall pigmentation. This region having a causal effect on pigmentation is bolstered by two additional results: (1) its ability to drive expression in the dermis of *Mus* embryos at a stage when relevant to the establishment of pigmentation prepatterns, and (2) patterns of DNA polymorphism show a strong signature of positive selection in this same small region. Interestingly, this region had not been previously identified as functionally important in *Mus* (Vrieling et al., 1994) or *Peromyscus* sp. (Linnen et al., 2013; Mallarino et al., 2017). Moreover, this region is not highly conserved (in rodents), suggesting it may have evolved regulatory function only recently. This new regulatory element further supports the observation that *Agouti* regulation is highly modular (Linnen et al., 2013; Mallarino et al., 2017; Vrieling et al., 1994), which could, in turn, explain why *Agouti* expression may be the target of repeated evolutionary tinkering across vertebrates, for example, in rabbits (Jones et al., 2018), dogs (Bannasch et al., 2021), buffalo (Liang et al., 2021) and birds (Nadeau et al., 2008; Uy et al., 2016) – akin to other highly modular genes, such as *Pax6* in vertebrates (Kammandel et al., 1999); *Pitx1* in stickleback fish (Chan et al., 2010; Thompson et al., 2018); *Ebony* in *Drosophila* (Signor et al., 2016); and *Asip2b*/*Agrp2* in cichlid fishes (Kratochwil et al., 2018).

This *Agouti* regulatory element likely contains causal mutation(s) that affect pigmentation. In total, there are 11 fixed differences between the light and dark *Agouti* alleles found in beach and mainland mice. While precisely which variant(s) are causal remains unclear, the three of these SNPs that are significantly associated with overall pigmentation (pPC1) in the polymorphic *albifrons* population represent the strongest candidates (Table S5). We also observe several complex indels and repetitive elements in the same region, which may themselves affect Agouti expression and drive an association signal in linked SNPs. While many of these variants disrupt predicted transcription factor binding sites identified in *Mus* (Table S4), such predictive approaches have poor specificity, and it is unclear if or when any of these sites may be active. And because this region is not well conserved even among rodents, results from gene-editing experiments in *Mus*, particularly ones that could result in subtle variation between mutants, may be challenging to interpret. However, surveying additional individuals in the admixed *albifrons* population may allow us to pinpoint the causal mutation(s) in the future. Similarly, the establishment of dermal cell lines that express Agouti, which are currently unavailable, containing the correct *trans* environment could allow us to test the effects of specific mutations – or combinations of mutations – via luciferase assays.

With a regulatory element identified, one can now more easily determine the source of this variation. In our survey of *Agouti* variation across both beach and mainland populations, we found the light *Agouti* allele is fixed in beach populations (see below), but also segregating or even fixed in some dark mainland populations. Beach mice are known to have very small populations (Oli et al., 2001; Mullen et al., 2009), thus the opportunity for a new adaptive mutation to arise is low (Domingues et al., 2012). Moreover, because the beach habitat is relatively young (8-10 kya; MacNeil, 1950) and colonization of that habitat relatively recent (Domingues et al., 2012), there has been only limited time for a new mutation to arise, and much less time for this mutation to spread to and across the mainland via migration. Thus, a more parsimonious scenario suggests that the light *Agouti* allele was selected from standing genetic variation existing in the mainland population(s), possibly at mutation-selection balance. This is consistent with results from *Mc1r*, in which the age estimate for the emergence of the causal *Mc1r* mutation predates the age of the beach habitat (Domingues et al., 2012). Interestingly, this scenario was predicted almost a century ago by Francis Sumner (1926) based on reports of light-colored mice occurring on the mainland near isolated beach habitats (Howell, 1920). Taken together, our data indicate that the large mainland populations are likely to have been the source of the light *Agouti* allele.

This evolutionary scenario then raises the question of how the light *Agouti* allele is maintained in mainland populations, where it may be deleterious. Indeed, previous field experiments demonstrated that models of light mice experienced higher rates of predation than dark models in a dark-soil mainland habitat (Vignieri et al., 2010). We discuss two possible, and non-mutually exclusive, explanations. First, many mainland mice that carry the light *Agouti* allele (even homozygotes) appear to have relatively dark coloration typical of a mainland mouse. In *Peromyscus*, *Agouti* is known to contain multiple mutations that affect pigmentation (Linnen et al., 2013) and to interact with other pigmentation genes (e.g., Steiner et al., 2007); therefore, it is possible that epistatic interactions between mutation(s) in the newly discovered regulatory element and other mutations in *Agouti* or elsewhere in the genome explain why, in some populations, the light *Agouti* allele has minimal effect on pigmentation (see Manceau et al., 2010), thus limiting its visibility to selection. Indeed, previous work in beach mice demonstrated a role for epistasis between *Mc1r*, *Agouti* and *Corin* (Steiner et al., 2007; Manceau et al., in review). Second, in some mainland populations, such as *P. p. albifrons*, soil coloration is not uniformly dark, but rather patchy, with sometimes large regions of surprisingly light, beach-like substrate. Such mainland areas have light sandy soil due to the geological history of the southeastern US, which has experienced successive episodes of glacial advance and retreat, depositing light sediments inland and forming sand-dune habitats that remain to this day (Lane, 1994). Thus, the light *Agouti* allele may, at least in some mainland populations, be beneficial, consistent with a signature of positive selection on the light allele in the *albifrons* population. However, the *P. p. albifrons* population also harbors dark *Agouti* alleles, possibly due to migration from surrounding dark mainland populations (Haldane 1948, Mullen and Hoekstra, 2008) or spatially variable selection in patchy habitats (Pfeifer et al., 2019). Additional sampling through the range of *P. polionotus,* including measurements of soil color, combined with whole-genome re-sequencing may shed further light on these two hypotheses to explain the prevalence of the light *Agouti* allele in mainland populations.

While the light *Agouti* allele is found at varying frequency in mainland populations, this allele is fixed in both Gulf and Atlantic coast beach mice. The distribution of the light allele is consistent with a scenario in which the same allele was independently selected in the two beach lineages from standing genetic variation. We note, however, that without specific information about the causal mutation(s), we cannot rule out the formal possibility that independent mutations with similar phenotypic effects evolved within the same small regulatory region. However, given that the Gulf and Atlantic clades are closely related and recently derived from similar mainland ancestors (Steiner et al., 2007), some may argue this was an ideal scenario for repeated selection on shared ancestral variation (e.g., Bolnick et al., 2018; Conte et al., 2012). Sharing the same *Agouti* light allele would provide a simple mechanistic explanation for why the Gulf and Atlantic Coast beach mice are so similar in coloration (see Hoekstra et al., 2006).

Together, these results suggest a scenario in which a *cis*-acting regulatory mutation(s) in *Agouti* likely evolved in the mainland and was independently selected in both the Gulf and Atlantic coast beach mice, contributing to their rapid, parallel evolution. This evolutionary history is in stark contrast with previous results for a second pigmentation gene *Mc1r* (Hoekstra et al., 2006). In this latter case, it is a coding change (i.e., a single amino-acid mutation that reduces receptor signaling) that contributes to light coloration in beach mice. Also, this *Mc1r* mutation is found in (some but not all) Gulf coast beach subspecies but is completely absent from Atlantic coast beach mice (Steiner et al., 2009), thereby not contributing to their parallel color adaptation. Thus, together, these two genes demonstrate how – even within a single species and associated with the same phenotype – evolution may take very different genetic paths to similar phenotypic ends.

## Acknowledgements

We thank M. Omura of the Harvard Museum of Comparative Zoology (MCZ) for assistance in preparing and accessioning voucher specimens, N. Bedford for collecting *P. p. leucocephalus* specimens for whole genome sequencing, and E. Kingsley for providing *P. m. nubiterrae* samples, J. Weber and E. Delaney for assistance in the field. C. Hu designed the illustration in Fig. 2A. T. Capellini provided advice on the reporter assays and feedback on a draft of this manuscript together with S. He, C. Kratochwil and M. Manceau. AFK was supported by postdoctoral fellowships from the European Molecular Biology Organization (EMBO; ALTF 47-2018) and the German Science Foundation (DFG; KA 5308/1-1), JML from EMBO (ALTF 379-2011), the Human Frontiers Science Program (LT001086/2012), and the Belgian American Educational Foundation, and VSD from the Portuguese Foundation for Science and Technology. Fieldwork was supported by a MCZ Putnam Grant (to VSD and HEH), and lab work by the NIH (R35GM133758 to RM) and NSF (DEB-0919190 to HEH). HEH is an Investigator of the Howard Hughes Medical Institute.

## Author contributions

VSD and HEH conducted the field collections. JML generated the genome assembly annotation, and VSD generated the sequence capture data. TBW, AFK, and JML generated whole genome resequencing data. TBW and AFK performed all additional genomic analyses. TBW and SM scored pigmentation traits. TBW and RM designed the reporter assays, and RM performed tissue sectioning. TBW, AFK, and HEH wrote the manuscript, with input from all authors.

## Materials and Methods

### Specimen and tissue collection

Over two expeditions, in the summer and winter of 2009, we collected 168 *Peromyscus polionotus albifrons* mice from a single population occupying habitat with patches of both light-colored sand and dark loam-clay soil in Lafayette Creek Wildlife Management Area of Walton County, Florida, approximately 25 km inland from the Gulf of Florida (Table S1). Mice were captured overnight using Sherman live traps. Following euthanasia, we sampled liver tissue from each individual and placed the tissue in 95% ethanol until they could be transferred to −80°C for long-term storage. We also prepared standard museum skins and skeletons. Both tissue and specimens were then accessioned in Harvard University’s Museum of Comparative Zoology (MCZ). In addition to *P. p. albifrons*, we included specimens from 10 distinct beach and mainland locations – representing eight additional *polionotus* subspecies – across the southeastern United States, as well as *P. maniculatus nubiterrae* from the northeastern US as an outgroup (Table S1). Tissues and voucher specimens are accessioned in the Harvard MCZ (Table S5).

### Measurement of pigment variation

We measured 23 pigmentation traits on specimens prepared as flat skins using two approaches: (1) the distribution of pigmentation (e.g., tail-stripe length) or (2) the intensity of pigment (e.g., dorsal hue, brightness). These specific traits were chosen because they are known to vary among beach mouse populations (see Mullen et al., 2009; Steiner et al., 2007; 2009); we did not find any new body regions that showed measurable variation with the *albifrons* population. We first scored the extent of pigmentation for 14 body regions, including dorsal, flank, and ventrum pigmentation, rump shape and rump shadow, ankle shadow, tail stripe, ear base, eyebrows, cheek, whiskers, rostrum, and between the eyes (see Fig. 2A). A test set of 10 individuals were scored by two independent researchers, and the methods refined until their scores were identical. For the full dataset, each trait was scored by a single individual across all specimens to ensure consistency: two researchers scored traits with each scoring half the traits in all individuals. Second, to measure pigment intensity, we used a FLAME UV-VIS spectrometer with a pulsed xenon light source, a 400 μm reflectance probe, and OceanView software (Ocean Optics) to measure five reflectance spectra from each of three body regions (dorsal stripe, flank, and ventrum). We used a custom *R* script to obtain brightness, hue, and saturation values in the visible spectrum (400-700 nm) with 1 nm bin width, using a segment classification approach (Endler, 1990) with formulae as described for CLR v 1.05 (Montgomerie, 2008). For all traits, we took five measurements and then calculated the median value for each body region for each individual. In total, we measured these 23 traits on 168 *P. p. albifrons* specimens as well as representative individuals from the Gulf (*P. p. leucocephalus*, n= 13), Atlantic (*P. p. niveiventris*, n= 15), and mainland (*P. p. polionotus*, n= 17) populations.

### Trait correlations and phenotypic PCA

To test for correlations among traits, we calculated pairwise trait correlations using the cor(., method=“pearson”, use=“complete.obs”) and cor.mtest() functions in *base R*, correcting for the number of pairwise tests to determine statistical significance (Bonferroni method). To account for trait correlations and to reduce the dimensionality of our dataset, we performed a principal component analysis (PCA) of all pigmentation traits using the *FactoMineR* (Lê et al., 2008) and *factoextra* (Kassambara and Mundt, 2020) *R* libraries. More specifically, we first estimated the best number of dimensions for imputing missing data with *estim_ncpPCA(., method.cv=“Kfold”)*, imputed missing data based on the estimated number with *imputePCA(., ncp=5)* and then performed the PCA on the imputed dataset using the *PCA()* function. Pairwise trait correlations and the phenotypic PCA were based on *P. p. albifrons* individuals only. To compare overall pigmentation (largely captured by phenotypic PC1) among populations, pigment scores for individuals from other populations were projected onto the *albifrons* principal component space post hoc using the *predict() R* function.

### Genome sequencing and assembly

DNA was extracted using standard laboratory procedures from the liver of one female (*Peromyscus polionotus subgriseus;* PO stock) obtained from our laboratory colony. By choosing a female individual, we have equal coverage for the autosomes and the X chromosome, but the Y chromosome is not part of the assembly. We prepared libraries with Illumina TruSeq DNA Sample Prep Kit v2 according to the manufacturer’s instructions, and performed *de novo* sequencing on an Illumina HiSeq platform using a combination of short paired-end libraries and longer mate-pair libraries suitable for use with the ALLPATHS-LG genome assembler (Gnerre et al., 2011). All libraries were constructed and sequenced at the Broad Institute Sequencing Platform (Cambridge, MA, USA). In total, we generated 240.27 Gb of raw sequence data, representing a total physical coverage of 290x, and assembled these reads using ALLPATHS-LG (version R48559).

We used ALLMAPS (Tang et al., 2015) in combination with five genetic maps based on interspecific crosses (RAD-based: Bendesky et al., 2017; Fisher et al., 2016; Kingsley 2015; Weber et al., 2013; gene-based: Kenney-Hunt et al., 2014) to assemble the scaffolds into the pseudo-chromosomes. DNA sequences corresponding to 182 genes and RAD markers used to build the genetic maps were aligned against the genome using BLAT (Kent, 2002). Markers that could not be unambiguously mapped to a single location in the genome were filtered out. A total of 58,922 markers were included in the dataset. During a first iteration, ALLMAPS (Tang et al., 2015) revealed that a total of 66 scaffolds housed markers associated with more than one linkage group and were likely mis-assembled. These were subsequently split, and the position of the breakpoints determined based on the ALLMAPS predictions and the location of discordantly mapped reads. In most cases, these corresponded to assembly gaps. After correcting for these assembly errors, ALLMAPS was ran an additional time to generate the pseudo-chromosomes. Our final assembly includes 531 scaffolds, encompassing 2,575,648,500 bp (97.4% of the total assembled sequence), distributed in 23 autosomes and the X chromosome. The orientation of 461 scaffolds corresponding to 2,566,039,849 bp (97.0% of the total sequence) could be determined due to the presence of more than one marker. We assigned chromosome names based on previous reports from interspecific reciprocal whole chromosome painting, which have allowed to assign linkage groups with known genes to *Peromyscus* chromosomes (Kenney-Hunt et al., 2014, Brown et al., 2018). The chosen chromosome assignments reflect the standardized *Peromyscus* cytogenetic nomenclature (Greenbaum et al., 1994).

### Genome annotation

We annotated repetitive elements using a combination of *RepeatModeler* (Smit and Hubley, 2008) and *RepeatMasker* (Smit et al., 2013) using *Peromyscus*- and rodent-specific repeat libraries. To annotate protein-coding genes, we used a recently developed annotation strategy making use of multiple genome alignments and an existing high-quality annotation set (Fiddes et al. 2018). While permitting the finding of newly discovered genes via *ab initio* gene modeling, this approach allows to identify orthology relationships readily and with high accuracy. We first aligned the oldfield mouse chromosome-level assembly to the assemblies of the laboratory mouse (*Mus musculus;* GRCm38), the rat (*Rattus norvegicus*; Rnor_6.0), the prairie vole (*Microtus ochrogaster;* MicOch1.0) and the prairie deer mouse (*Peromyscus maniculatus;* Pman2.1.3) using ProgressiveCactus (Paten et al. 2011a, 2011b). We reasoned that including more species that represent progressive level of evolutionary divergence would improve the accuracy of the ancestral sequence reconstruction process that takes place during the preparation of the whole-genome alignment. Using the Comparative Annotation Toolkit (CAT; Fiddes et al., 2018), we annotated the oldfield mouse genome using the genome of *Mus musculus* (GRCm38/mm10) and the high-quality and well-curated GENCODE VM15 as the reference gene/transcript set, as well as extensive transcriptome sequencing datasets for *P. polionotus* corresponding to five tissues (brain, testis, hypothalamus, main olfactory epithelium and vomeronasal organ).as well as skin RNA-Seq data from the prairie deer mouse, *P. maniculatus bairdii*.

To obtain quantitative measures of the completeness of the genome assembly, we used BUSCO (Simão et al. 2015; version 3.0.2) with BLAST+ (version 2.2.28+), HMMER (version 3.1b2) and AUGUSTUS (version 3.3.2). We used *human* as species, which specifies the parameters used by AUGUSTUS, and the *mammalia* and *euarchontoglires* sets for our analyses.

### Population sequencing, variant calling, and genotype likelihoods

For high coverage whole-genome sequencing (WGS) of representative beach (*P. p. leucocephalus*) and mainland (*P. p. subgriseus*) populations, we extracted DNA from ~20mg of liver tissue and generated sequencing libraries using a PCR-free KAPA HTP kit. Following enzymatic fragmentation, we used size selection to enrich for a 450bp insert size and ligated Illumina adapters. We sequenced the resulting libraries using 150bp paired-end sequencing on an Illumina NovaSeq S4 flowcell to achieve 15-20X coverage.

For additional Gulf, Atlantic, and mainland populations, we used a sequence capture strategy aimed at sequencing both (1) putatively neutral loci and (2) the pigmentation genes *Agouti* and *Mc1r*. Specifically, we targeted ~5000 1.5 kb non-coding regions randomly distributed across the genome as well as 190 kb and 150 kb regions flanking the *Agouti* and *Mc1r* loci, respectively (see Domingues et al. 2012 for capture array design details; Kingsley et al., 2009 for *Agouti* and *Mc1r* sequencing details). This strategy was applied to five Gulf beach mouse subspecies (*P. p. ammobates, P. p. allophrys*, *P. p. trisyllepsis, P. p. peninsularis, P. p. leucocephalus*), three mainland subspecies (*P. p. polionotus, P. p. albifrons,* three populations of *P. p. subgriseus*) and one Atlantic beach subspecies (*P. p. niveiventris*) (see Table S1). The availability of both high-quality WGS and sequence capture data for the *P. p. leucocephalus* and *P. p. subgriseus* subspecies allowed us to verify that the sequence capture loci accurately represented each population’s genetic diversity.

For both WGS and sequence capture data, we converted raw fastq files to unmapped bam files using *FastqToSam* (*Picard* toolkit, 2019) and then marked Illumina adapters using *MarkIlluminaAdapters* (*Picard*). Using *SamToFastq* (*Picard*), we created interleaved fastq files and clipped adapter sequences. We mapped sequencing reads to the *P. polionotus subgriseus* reference genome (see above) using *bwa-mem* (Li and Durbin, 2009), with –p to indicate interleaved paired-end fastq input, and –M to mark short split hits as secondary for compatibility with Picard. We then used *MergeBamAlignment* (*Picard*) to merge mapped and unmapped bam files to preserve read group information and sequencing duplicates using *MarkDuplicates* (*Picard*), with OPTICAL_DUPLICATE_PIXEL_DISTANCE=2500 to account for artifacts generated from the patterned flowcell found in the NovaSeq S4.

We then called variants separately for the WGS and sequence-capture datasets to reduce processing time, as they vary significantly in both coverage and sample number. However, the following variant calling and filtering steps were applied equally to both data types. To begin, we used *HaplotypeCaller* (GATK 3.8; Poplin et al., 2018) on the aligned bam files with the default heterozygosity prior (-hets = 0.005) and –ERC GVCF to produce per-sample gVCFs. For the X chromosome, we specified a prior input ploidy based on a comparison of coverage with the autosomes using *samtools* (v. 1.10) *depth* (Li et al., 2009). Next, for the WGS data, we generated variant + invariant cohort-level vcfs for each chromosome using *GenotypeGVCFs* (*GATK 3.8*) with “--max-alternate--alleles 4 -all-sites”. For the sequence capture data, the “-allsites” parameter was removed and only variants were reported. These raw, cohort level vcfs were split into indels and SNPs with *SplitVcfs* (*Picard*) and invariant sites with *SelectVariants* (*GATK 3.8*). We performed filtering on each set independently, excluding SNPs with QD < 2.0, FS > 10.0, MQ < 40.0, MQRankSum < −12.5, ReadPosRankSum < −8.0 or SOR > 3.0 and excluding INDELs with QD < 2.0, FS > 200.0, ReadPosRankSum < −20.0, SOR > 3.0. We also retained invariant sites with QUAL ≥ 20 using *bcftools 1.11-95* (Li, 2011). These filtering parameters were based on a combination of GATK recommendations for datasets without truth/training sets, and visual inspection of the distributions for each metric. We also set individual genotype calls to missing if the read depth at a given site was less than five. Finally, we combined the sequence capture dataset with the WGS dataset using *vcf-merge* (*vcftools* 0.1.15; Danecek et al., 2011).

### Estimation of population structure

To test for population structure, we ran a genetic principal components analysis (gPCA) using *PCAangsd*, which is specialized for use with low-coverage, high-throughput sequencing data (Meisner and Albrechtsen, 2018). We used *beagle* genotype probability files for all sequence-capture loci as input and ran the program with default parameters. Using the output covariance matrix, we calculated eigenvalues and eigenvectors with the base R function *eigen*. We estimated population differentiation (Fst) for all pairwise population comparisons using the program *ANGSD* (Korneliussen et al., 2014). We first calculated the two-population site frequency spectra (2DSFS) using the SAF files generated by *ANGSD*, running *realSFS* with default parameters. We then generated the Fst index for each population pair with *realSFS fst index*, supplying each population’s SAF index and the 2DSFS with default parameters. The resulting Fst index file allowed us to estimate global Fst as well as Fst in sliding windows, using *realSFS fst stats* and *realSFS fst stats2*, respectively.

### Estimation of population relationships

To estimate the relationships among the sampled subspecies, we constructed a population-level tree using a modified version of *SNAPP*, a multispecies coalescent-based tool that uses biallelic markers as input (Bouckaert et al., 2019; Bryant et al., 2012; Stange et al., 2018). Our input data consisted of genome-wide putatively neutral variants sampled in both the sequence-capture and whole-genome datasets (i.e., excluding the *Agouti* and *Mc1r* regions). Briefly, we chose the two highest-coverage individuals representing each population, then retained biallelic SNPs with minor allele frequency greater than 0.05, excluded variants that violated Hardy-Weinberg equilibrium (p-value < 0.001) in four or more populations, and thinned the remaining variants so that none were within 100 bp of each other. The remaining variants were reformatted as a phylip file and converted to the xml format required by *SNAPP/BEAST* using the script *snapp_prep.rb* (https://raw.githubusercontent.com/mmatschiner/snapp_prep/master/snapp_prep.rb). To specify a starting tree constraint (-s), we ran *RAxML* (Stamatakis, 2014) with ascertainment bias correction (--asc-corr=lewis) on a reduced dataset containing the highest coverage representative of each subspecies to obtain a maximum likelihood phylogeny. We also specified a node constraint (-c) that the crown divergence of all subspecies, excluding *P. maniculatus nubiterrae* (outgroup), should approximate a normal distribution mean of 8.9 kya and a standard deviation of 1.5 kya. These values were taken from *SMC++* estimates of the divergence time between mainland (*P. p. subgriseus*) and beach (*P. p. leucocephalus*) subspecies, assuming a generation time of four months (i.e., 3 generations/year; see ‘Demographic Inference’ below). Finally, we sampled 1000 random variants from the remaining dataset to speed up run times and specified 1 million MCMC iterations. For quality control, we confirmed thorough mixing in the run using *Tracer v1.7.1* (Rambaut et al., 2018) and visually inspected the trees using *DensiTree* (Bouckaert, 2010). A consensus tree was generated with *TreeAnnotator* (Bouckaert et al., 2019), using a 10% burn-in and reporting mean node heights.

### Demographic inference

The whole-genome, high-density sequencing coverage for one mainland (*P. p. subgriseus*) and one beach (*P. p. leucocephalus)* subspecies allowed us to infer demographic histories with high resolution. Specifically, we used the program *SMC++* (Terhorst et al., 2016) to estimate population divergence times and parameterize population size changes in additional populations. To mask low-quality regions, we followed the *SNPable* protocol (http://lh3lh3.users.sourceforge.net/snpable.shtml) to identify regions in the assembly with poor mappability, using a k-mer size of 150 bp. SNPs that violated Hardy-Weinberg equilibrium (p < 0.01) and had low population coverage (<80% samples genotyped) were also excluded.

We then used the *vcf2smc* command to create the per-population SMC input files, supplying mappability, missingness, and Hardy-Weinberg masks to exclude low quality regions in the dataset. The ‘distinguished individual’ (DI), a key feature of *SMC++*, was specified as the highest coverage sample for each population. We generated two-population input files using the same command and input files, but with no specified DI (not applicable to multi-population analysis). For single-population inference, we used *cv* with the following parameters: ‘--folds 4 --timepoints 1e3 5e7 --Nmax 1e8 –spline cubic’ and a germline mutation rate of 5.3e-9 (Uchimura et al., 2015). We also ran *estimate*, an earlier version of *smc++ cv*, with identical parameters for downstream compatibility with population-split inference. We then provided single-population demographic models (as obtained by *smc++ estimate*) and two-population input files to *split* to estimate the timing of the mainland (*subgriseus*) and beach (*leucocephalus*) split. To obtain confidence intervals for all the estimates described above, we used a custom script to resample 10 Mb stretches of the genome in the SMC input files, thus generating 20 bootstrap replicates per estimate. The above pipeline was rerun with identical parameters on these replicates, and 95% confidence intervals were calculated as mean +− 2*standard error.

### Genome-wide association mapping

Genotype-phenotype associations were determined using the mixed-model approach implemented in *EMMAX (beta-07Mar2010)*, accounting for population structure / relatedness by incorporating a Balding-Nichols kinship matrix as a random effect (Kang et al., 2010). We set the statistical significance threshold at p<0.05 after correcting (Bonferroni method) for the number of effective independent tests. We obtained the latter using *Genetic Type I error calculator (GEC) v0.2* (Li et al., 2012). We excluded samples with more than 50% missing genotypes from these analyses, leaving N=152 samples. We used both biallelic SNPs and indels for association mapping, but excluded markers with > 50% missing data, a minor allele frequency < 0.05, or deviating from Hardy-Weinberg equilibrium (p<0.001). We generated Manhattan plots and QQ plots using the *qqman* (Turner, 2018) and *snpStats* (Clayton, 2021) *R* libraries, respectively. Using *plink* (v1.90b6.15), we calculated pairwise linkage disequilibrium (r^2^) among SNPs in the focal region (flags: --chr chr4 --from-bp 9820301 --to-bp 9870301 --r2 --ld-window-r2 0 --ld-window 1000). Next, we estimated the proportion of variance explained (PVE) for a given SNP (assuming Hardy-Weinberg equilibrium) using genotype-phenotype regressions.

### Sequence conservation

To evaluate the nucleotide sequence conservation level of the *Agouti* locus in *P. polionotus*, including the candidate regulatory region, we downloaded all available orthologous rodent *Agouti* sequences from NCBI (accessed pt. 7, 2020) using *esearch* (-db gene -query “ortholog_gene_434[group] AND rodents[orgn]”) in combination with *esummary* and *xtract* of the entrez direct e-utilities. Next, we manually added 15 kb to each of the start and end coordinates (or the maximum number of bp if hitting a scaffold end) using a custom *awk* script and retrieved the corresponding nucleotide sequences with *efetch*. The sequence of *Nannospalax galili* was removed due to a lack of available flanking sequence. Finally, we determined sequence conservation between *P. polionotus* and the remaining 26 rodent species using *mVISTA* (Frazer et al., 2004).

### Regulatory database queries

To determine if the candidate regulatory region of *Agouti* contains any known regulatory elements or transcription factor binding sites, we downloaded both phastCons60way conserved elements and ORegAnno regulatory elements from the UCSC genome browser in *Mus musculus* mm10 coordinates (http://hgdownload.cse.ucsc.edu/goldenpath/mm10/database/). Elements from each database were converted to *P. polionotus* genomic coordinates using UCSC’s *liftOver* utility (Kent et al., 2002) and a custom chain file, with the parameters ‘-multiple -minMatch=0.70’.

We obtained ENSEMBL regulatory features using the *R* package *biomaRt* (Durinck et al., 2009, 2005). The mm39 regulatory feature dataset was retrieved with the function *useDataset()*, with the parameters ‘dataset=“mmusculus_regulatory_feature”, mart= “ENSEMBL_MART_FUNCGEN”’, and *getBM()* was used to retrieve entries from the broader *Agouti* region using the extended *Agouti* coordinates (2:154785921:155055915) for the mm39 assembly. We directly converted coordinates in mm39 to mm10 assembly coordinates using *liftOver* with default parameters and the UCSC mm39toMm10 chain file (https://hgdownload.soe.ucsc.edu/goldenPath/mm39/liftOver/mm39ToMm10.over.chain.gz), then converted mm10 coordinates to *P. polionotus* coordinates using the same approach described above.

### LacZ reporter assay

To determine if the candidate region was capable of regulatory activity, we assessed whether it could drive expression of the *lacZ* reporter gene in the skin of developing mouse embryos (strain: FVB/NJ). To identify the most appropriate sequence length for this experiment, we specified boundaries that encompassed the three pPC1-significant SNPs, the unique local topology regions identified by *Saguaro* (see Methods: Local tree inference with *Saguaro*) and the tract of relatively high sequence conservation at the 3’ end of the association & *Saguaro* regions, resulting in a total sequence length of 2.6 kb (Fig. 4B). While *lacZ* experiments are particularly useful for verifying that a regulatory locus is active, comparisons between alleles of the same locus (e.g., light and dark alleles) can be challenging due to the noise associated with random genomic integration of the construct. Therefore, presented with two alternative haplotypes in this region – “light” and “dark” – we decided to use the light haplotype for these experiments, under the assumption that the light allele was less likely to contain mutations reducing element activity (i.e., high Agouti expression is generally associated with light pigmentation).

We used the *lacZ* expression vector, hsp68lacZ (gift from T. Capellini, Addgene #37843). The light haplotype sequence file and hsp68lacZ vector were provided to Taconic Biosciences (NY, USA) who synthesized and cloned the sequence upstream of the hsp68 minimal promoter, followed by pronuclear microinjection, collection of E14.5 embryos, genotyping, and *lacZ* staining. Stained embryos were shipped to our laboratory, where they were photographed, embedded in OCT, cryosectioned, and imaged.

### Transcription factor binding site prediction

To determine if variation in the regulatory element could be modifying relevant transcription factor (TF) binding sites, we examined motif differences at variant positions across the region. Specifically, we obtained all polymorphic sites in the regulatory element (chr4:9,844,852 bp – 9,847,500 bp) with minor allele frequency (MAF) > 0.05 in the *P. p. albifrons* population. We extracted the region 15 bp upstream and downstream of each variant (~30 bp sequence) and used *vcf-consensus* (*vcftools* 0.1.15) to create an alternate sequence incorporating the variant. For each reference and alternate sequence, we used *CiiiDER* v.0.9 (Gearing et al., 2019) to predict TF binding sites, providing the database of 251 *Mus musculus* CORE TF position weight matrices available on *JASPAR* (downloaded 10-20-21; Fornes et al., 2020).

### Haplotype homozygosity tests

To test for evidence of non-neutral evolution in patterns of nucleotide variation, we calculated haplotype statistics. We first ran *fastPHASE* to create phased variant calls (Scheet and Stephens, 2006). We converted the input vcf for all individual genotypes at *Agouti*, *Mc1r*, and the sequence-capture loci to the *fastPHASE* format with *vcf2fastPHASE.pl* (https://github.com/lstevison/vcf-conversion-tools), then ran *fastPHASE* with the following parameters: −T20 −H50 -F. We next converted the phased output back to the vcf format with *fastPHASE2VCF.pl*.

Using the *R* package *REHH*, we ran a series haplotype-based tests to scan for signatures of positive selection on the light and dark haplotypes (Gautier et al., 2017). For each population, we converted vcfs to an REHH-compatible file with a custom script (*hap2rehh.py*), using the *P. polionotus subgriseus* reference genome to polarize alleles. We then converted these files to haplohh objects with *data2haplohh*, then computed Extended haplotype homozygosity (EHH) statistics with *scan_hh*, with the parameters ‘discard_integration_at_border=FALSE, maxgap=2000’ to accommodate the sequence-capture dataset. We then ran *ihh2ihs* to calculate integrated haplotypes scores (IHS), using a minor allele frequency (MAF) filter of 0.05 and default allele frequency bin sizes of 0.025.

### Local tree inference with *Saguaro*

As a complementary approach to test for evidence of selection within *Agouti*, we used the Hidden Markov Model (HMM)-based software *Saguaro* to build local phylogenies from sequence data (Zamani et al., 2013). As input, we used variant calls from the sequence-capture dataset and the *Agouti* and *Mc1r* extended loci, and filtered out variants with MAF < 0.025. To reduce computational complexity and help with downstream interpretation, we reduced the sample size to include only the two highest-coverage representatives of each population. In the case of *albifrons*, we included two individuals homozygous for the “dark” allele and two for the ‘light’ allele (as determined by their genotype at SNP chr4:9,845,301bp). We then used *VCF2HMMFeature* to transform the variant calls to a *Saguaro*-compatible input format. We ran *Saguaro* for 15 iterations with default parameters. We transformed the resulting topologies to phylip files with *Saguaro2Phylip* and used a custom script to parse the *LocalTrees.out* file to obtain HMM transitions across the dataset.

---

**Table S1.** Sampling information. List of species, subspecies and populations included in this study. “Collector” points to information for the precise sampling method, time, and place. Primary collector initials are as follows: EPK = Evan Kingsley, NLB = Nicole Bedford, VSD = Vera Domingues. “Sequencing strategy” refers to either WGS = whole genome sequencing (highlighted in grey) or Seqcapture = targeted sequence capture array.

| Taxonomy | Collectors/<br>Publications | Latitude | Longitude | No. samples | Sequencing<br>strategy |
| --- | --- | --- | --- | --- | --- |
| *P. polionotus ammobates* | Mullen *et al.* 2009 | 30.229978 | -87.814703 | 15 | Seqcapture |
| *P. polionotus allophrys* | Mullen *et al.* 2009 | 30.077795 | -85.647814 | 11 | Seqcapture |
| *P. polionotus leucocephalus* | NLB | 30.397536 | -86.729057 | 15 | WGS |
| *P. polionotus leucocephalus* | Mullen *et al.* 2009 | 30.397536 | -86.729057 | 20 | Seqcapture |
| *P. maniculatus nubiterrae* | EPK | 40.33 | -79.27 | 1 | Seqcapture |
| *P. polionotus trisyllepsis* | Mullen *et al.* 2009 | 30.29371 | -87.463557 | 5 | Seqcapture |
| *P. polionotus subgriseus* | VSD | 29.1828333 | -81.795 | 15 | WGS |
| *P. polionotus albifrons* | VSD | 30.5411 | -86.075717 | 168 | Seqcapture |
| *P. polionotus* | Domingues *et al.* 2012 | 31.995717 | -85.082967 | 6 | Seqcapture |
| *P. polionotus subgriseus* (A) | Domingues *et al.* 2012 | 30.814577 | -84.954529 | 5 | Seqcapture |
| *P. polionotus subgriseus* (T) | Domingues *et al.* 2012 | 31.6459 | -84.225 | 5 | Seqcapture |
| *P. polionotus subgriseus* (O) | Domingues *et al.* 2012 | 29.207927 | -81.740378 | 5 | Seqcapture |
| *P. polionotus peninsularis* | Mullen *et al.* 2009 | 29.957198 | -85.462412 | 19 | Seqcapture |
| *P. polionotus niveiventris* | Steiner *et al.* 2009 | 27.923493 | -80.488186 | 4 | Seqcapture |

**Table S2.**
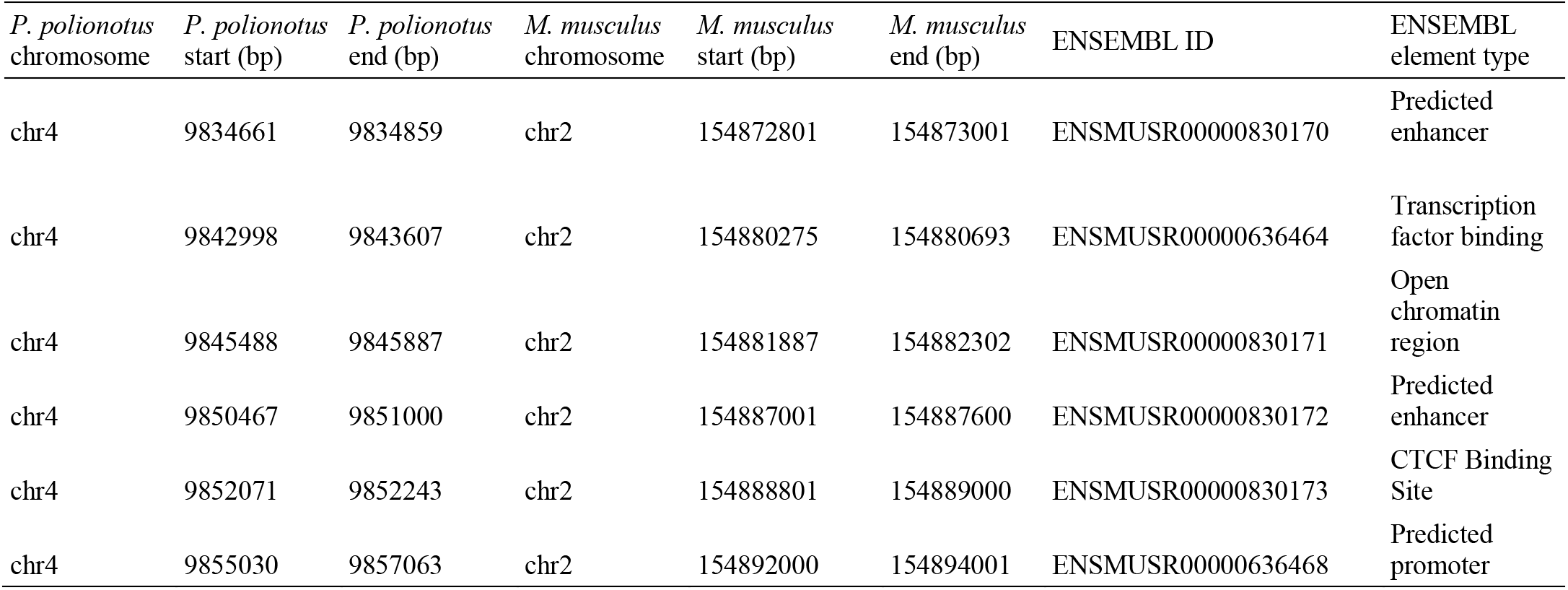
Regulatory elements in *Agouti*. *P. polionotus*-based coordinates (HU_Ppol1.3.3) of regulatory features lifted over from the ENSEMBL *Mus musculus* genome (GRCm39). Features found ~10 kb upstream and downstream of the 2 kb candidate region are shown.

| *P. polionotus*<br>chromosome | *P. polionotus*<br>start (bp) | *P. polionotus*<br>end (bp) | *M. musculus*<br>chromosome | *M. musculus*<br>start (bp) | *M. musculus*<br>end (bp) | ENSEMBL ID | ENSEMBL<br>element type |
| --- | --- | --- | --- | --- | --- | --- | --- |
| chr4 | 9834661 | 9834859 | chr2 | 154872801 | 154873001 | ENSMUSR00000830170 | Predicted<br>enhancer |
| chr4 | 9842998 | 9843607 | chr2 | 154880275 | 154880693 | ENSMUSR00000636464 | Transcription<br>factor binding |
| chr4 | 9845488 | 9845887 | chr2 | 154881887 | 154882302 | ENSMUSR00000830171 | Open<br>chromatin<br>region |
| chr4 | 9850467 | 9851000 | chr2 | 154887001 | 154887600 | ENSMUSR00000830172 | Predicted<br>enhancer |
| chr4 | 9852071 | 9852243 | chr2 | 154888801 | 154889000 | ENSMUSR00000830173 | CTCF Binding<br>Site |
| chr4 | 9855030 | 9857063 | chr2 | 154892000 | 154894001 | ENSMUSR00000636468 | Predicted<br>promoter |

**Table S3.**
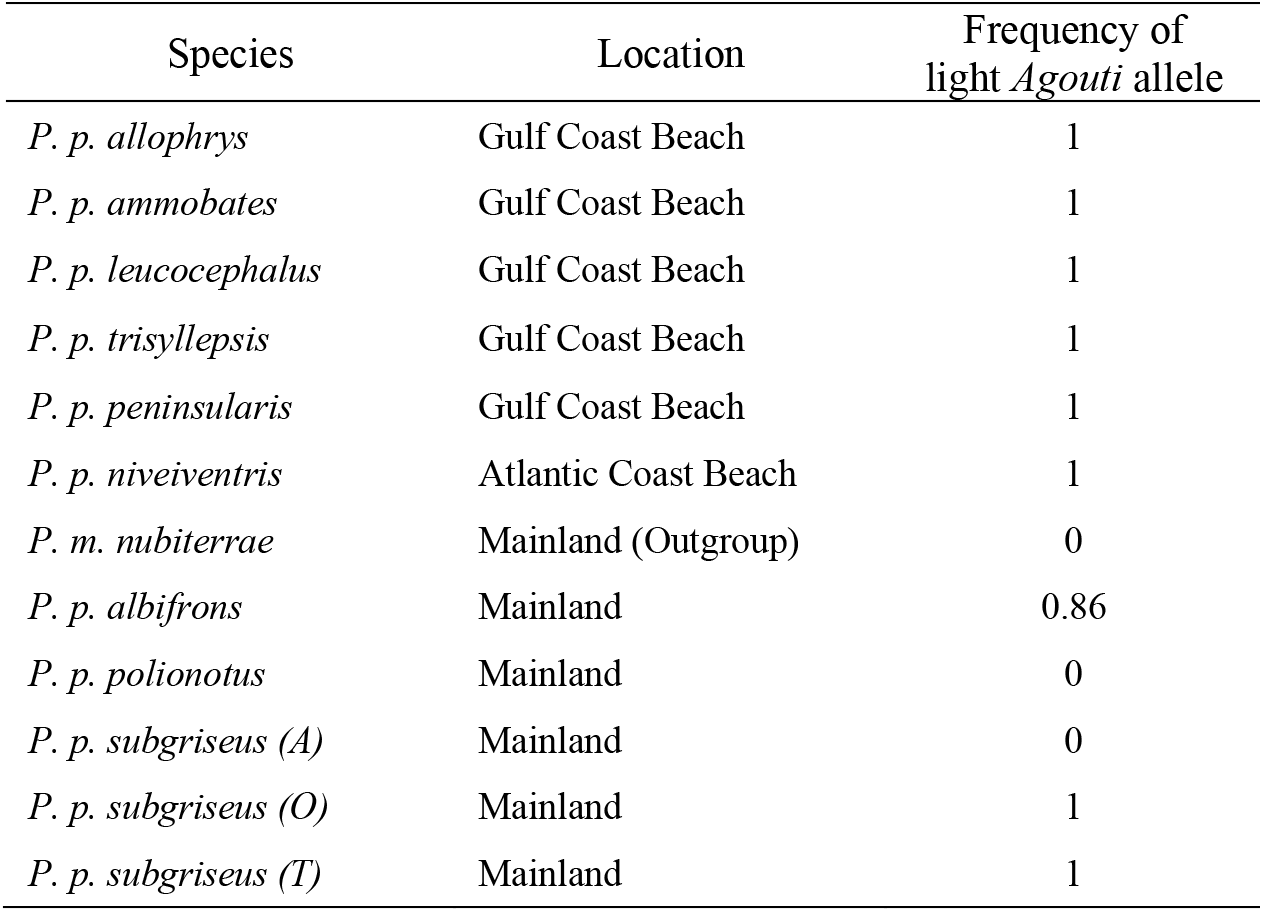
Population-level frequency of light allele. Each species is provided with its source location. Frequency of the light allele is calculated as the proportion of haplotypes in a population that have the light-associated allele at chr4:9,845,301 bp.

**Table S4.**
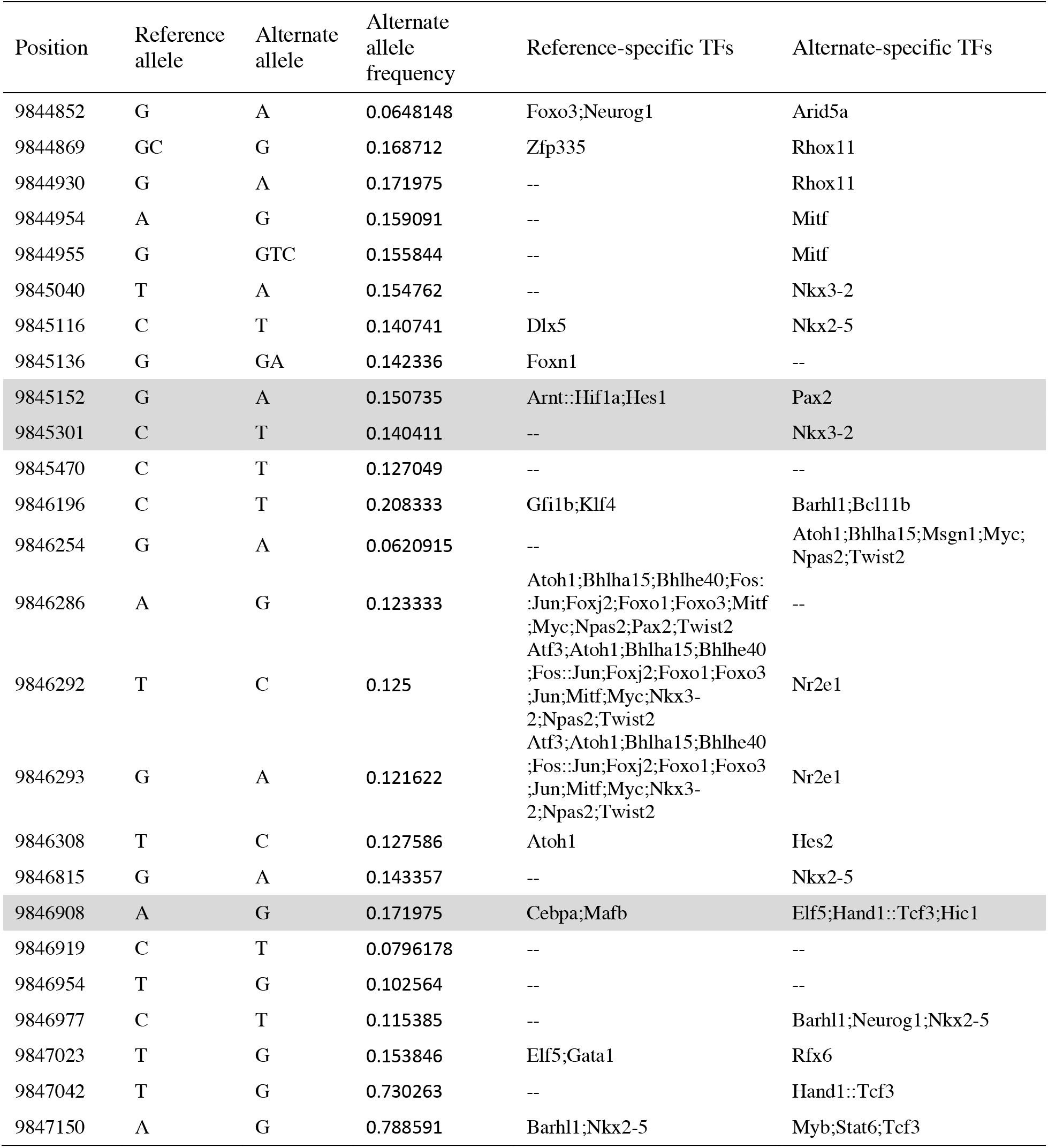
Transcription factor binding sites overlapping variant positions in the regulatory element of *P. p. albifrons*. “Position” includes only sites on chr4 in the tested regulatory element (9,844,852 bp – 9,847,500 bp) that are variant in *P. p. albifrons* with a minor allele frequency (MAF) > 0.05. Both “Reference allele” and “Alternate allele” are relative to the *P. p. subgriseus* reference genome (no relation to light or dark haplotypes). “Reference-specific TFs” and “Alternate-specific TFs” refer to predicted TF binding sites from JASPAR (see Methods) that are intact in the Reference allele and Alternate allele, respectively. pPC1-associated SNPs are highlighted in gray.

**Table S5.**
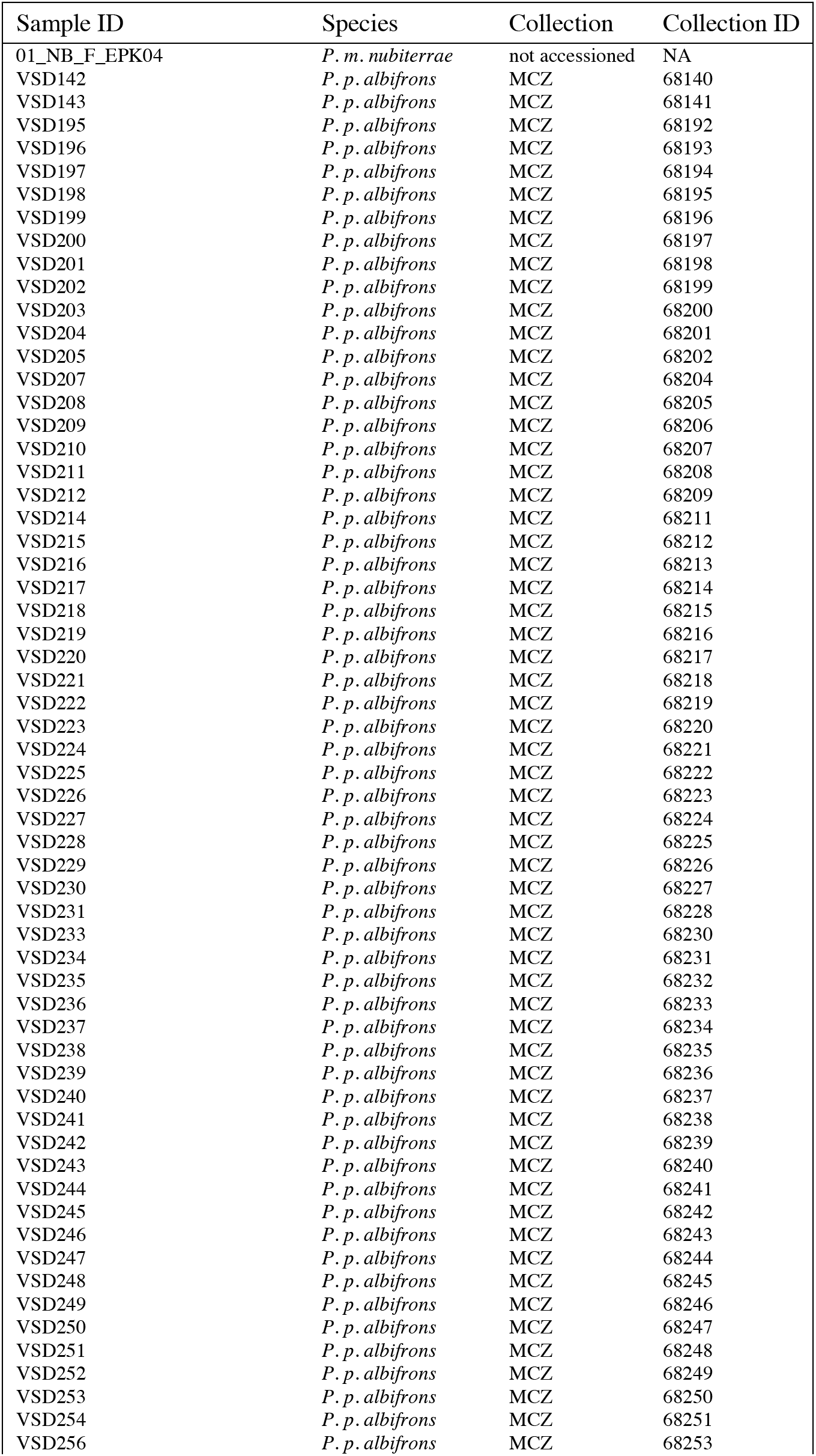
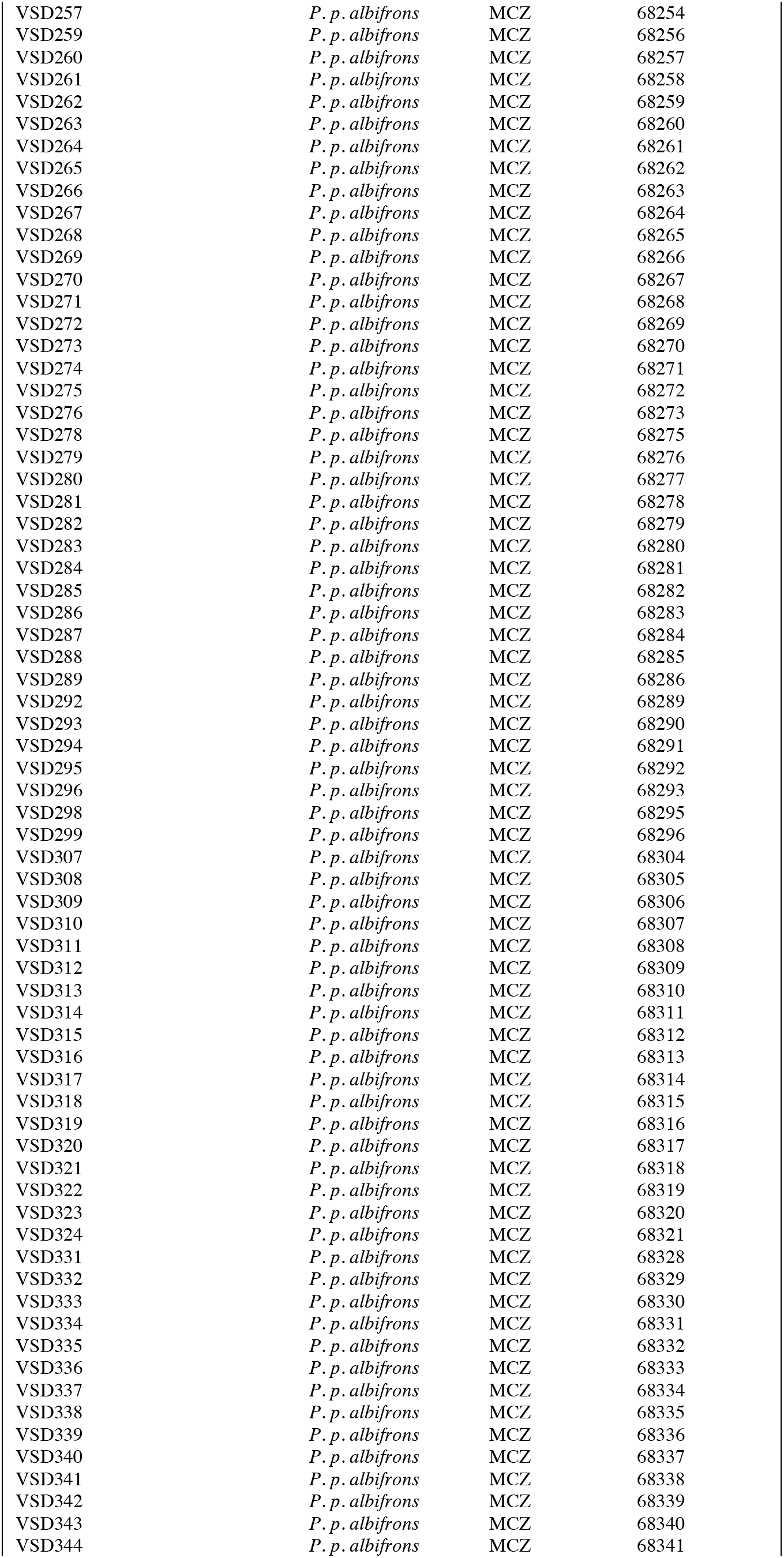
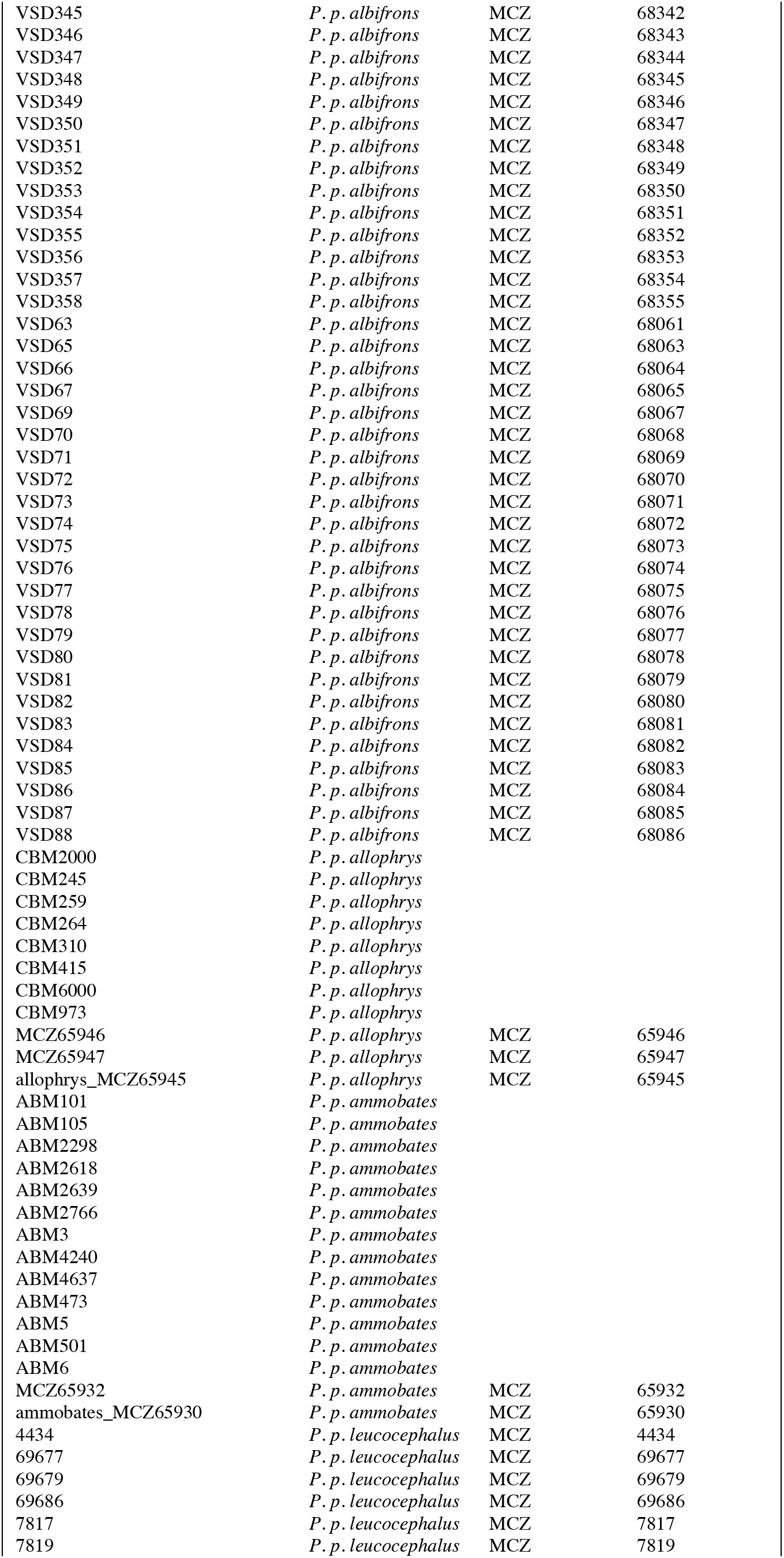
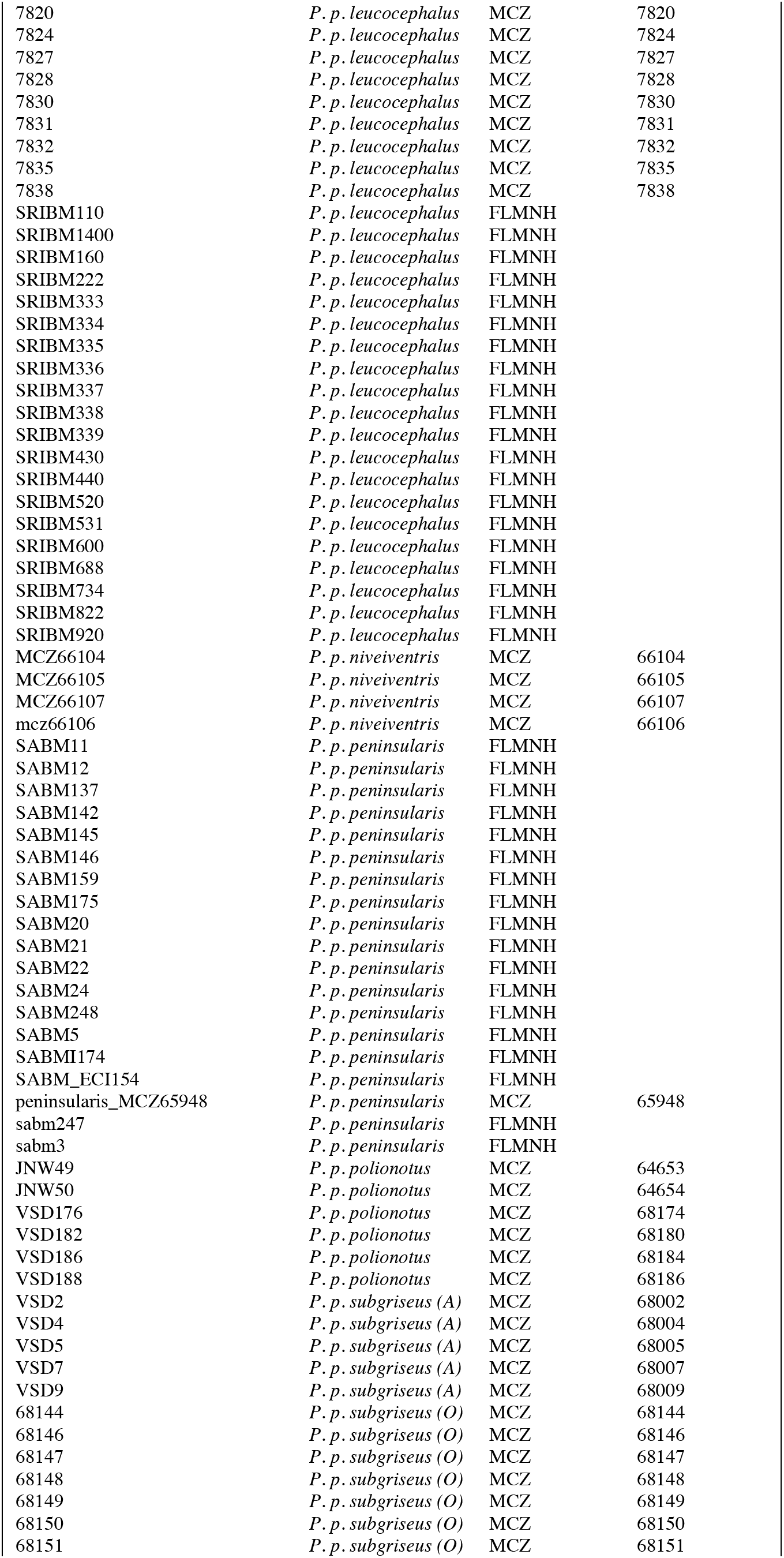
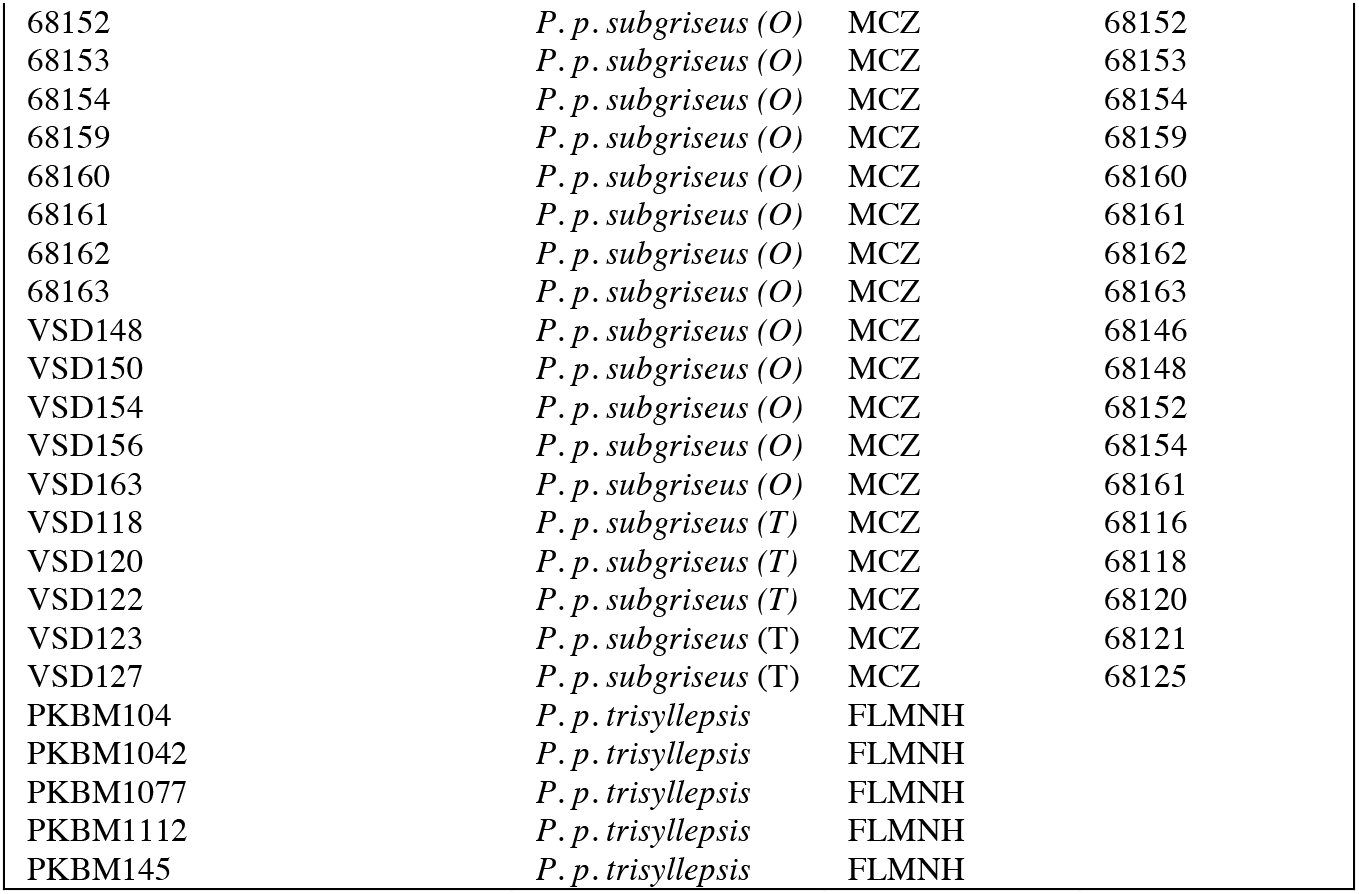
Museum accession information for all samples included in study. “Sample ID” refers to ID used by authors in this study, while “Collection ID” refers to the ID used by the institution where samples are accessioned. “Collection” refers to either MCZ = Harvard Museum of Comparative Zoology, or FLMNH = Florida Museum of Natural History.

**Figure S1.**
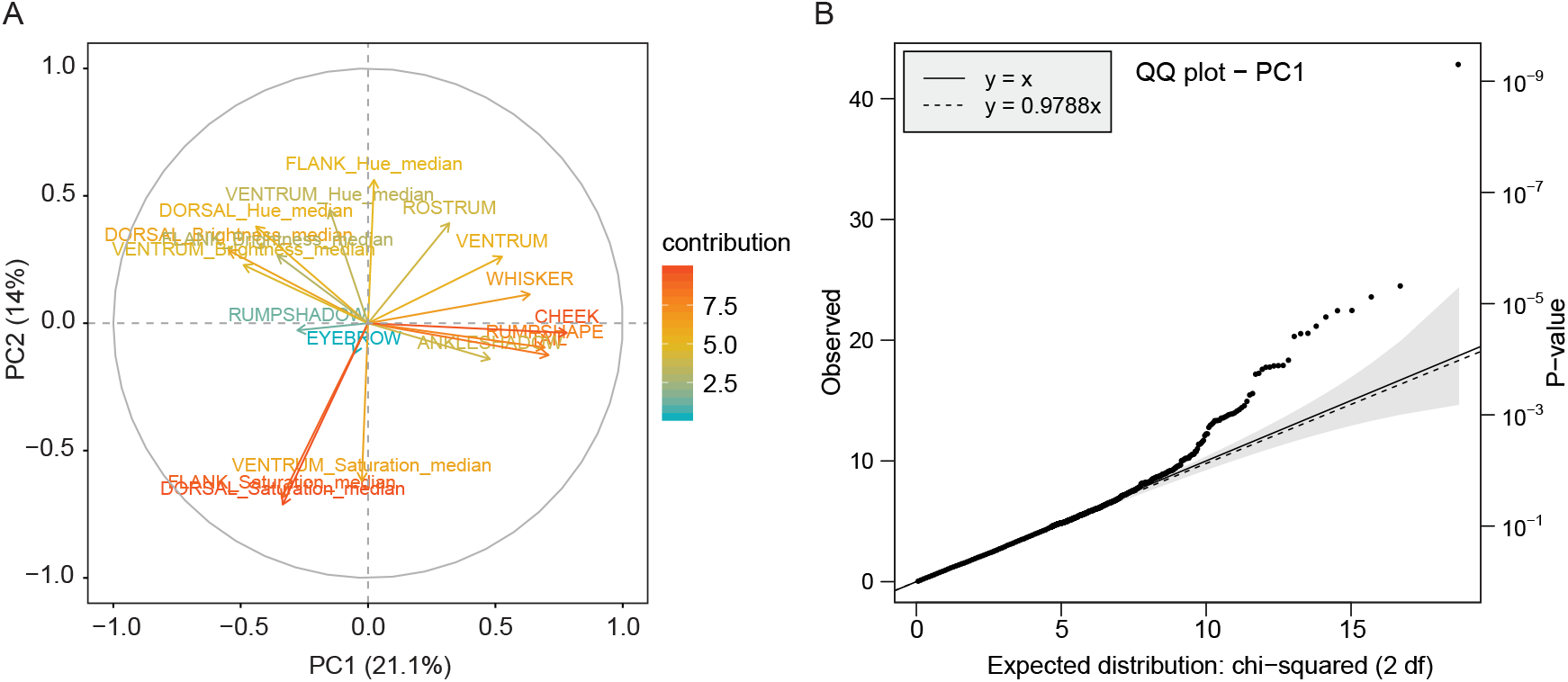
Pigment trait loadings on phenotypic Principal Component Analysis (PCA). **A**. PCA biplot shows contributions of pigment traits to the first two phenotypic PCs. Percentage values in parentheses correspond to the percent variance explained by each PC. **B**. Quantile-quantile plot of empirical vs. expected GWAS p-value distributions, indicating no signs of overdispersion or abnormal behavior.

**Figure S2.**
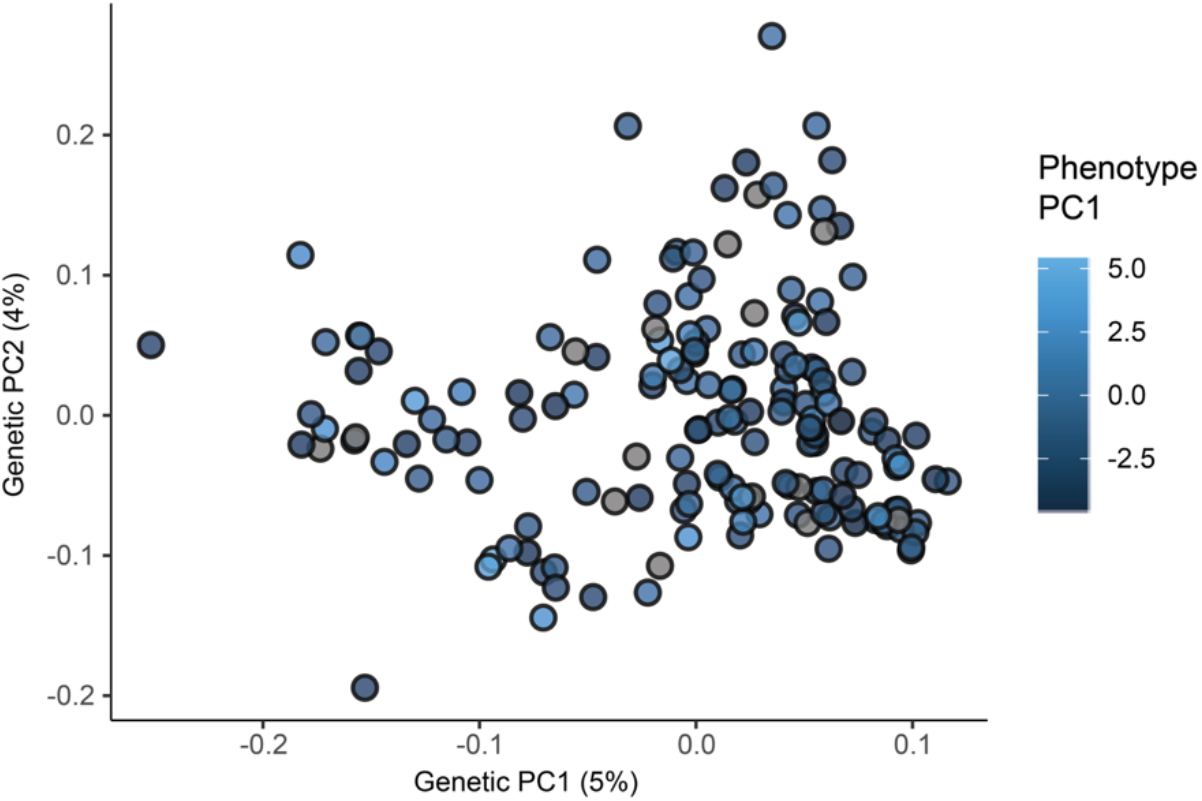
Genetic principal components analysis of the *P. p. albifrons* population. Each dot represents an individual (N=168). Color approximates phenotypic PC1 value. Percentage value on each axis corresponds to the percent variance explained by each genetic PC.

**Figure S3.**
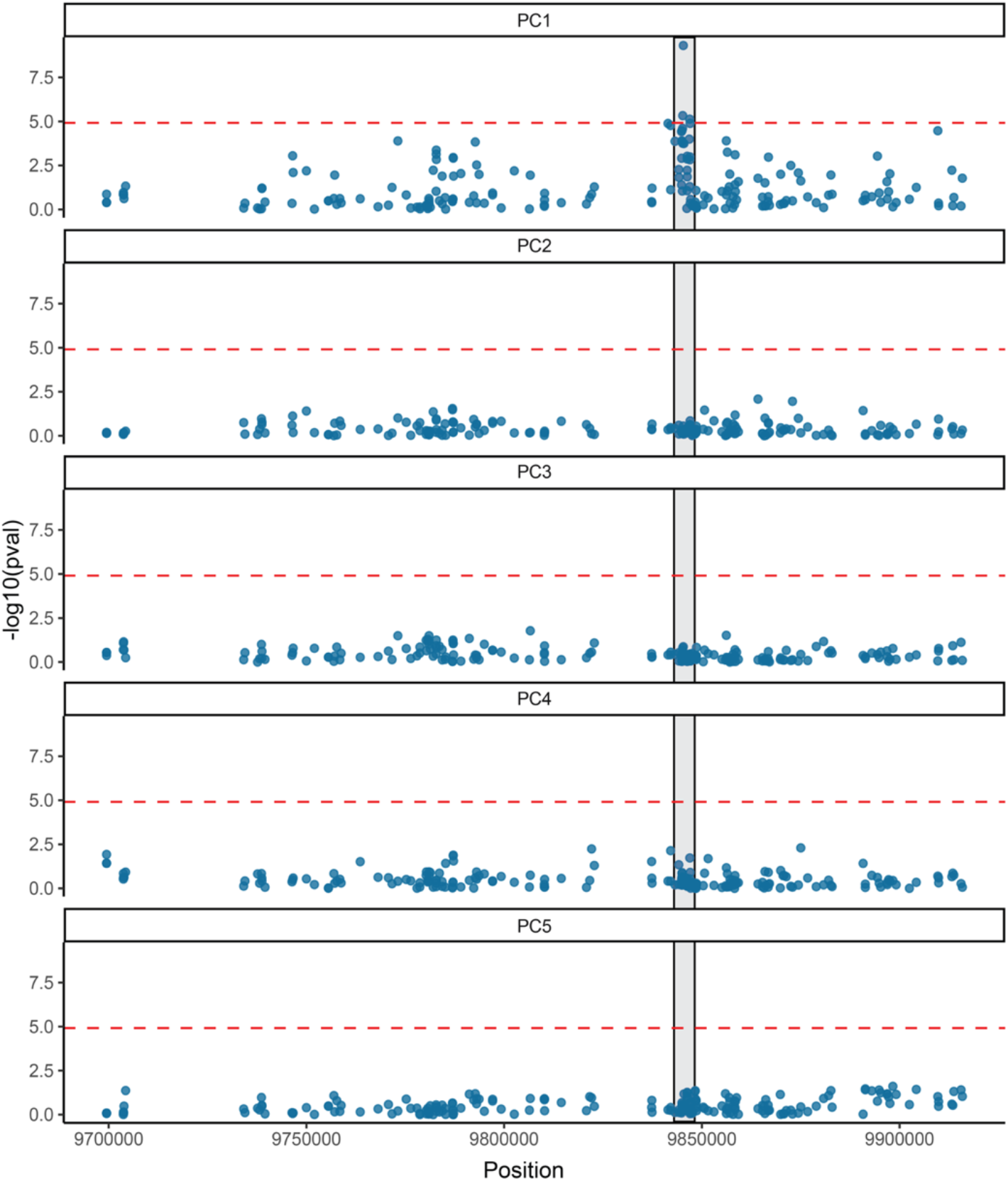
Manhattan plots showing association between phenotype and variation across the *Agouti* locus. Data for phenotypic PCs (pPCs) 1 to 5 are shown. Dashed red lines indicate genome-wide significant threshold, corrected for number of independent tests (see Methods). Gray bar denotes boundaries of peak association found for pPC1. No other pPCs show a significant association with variants in *Agouti.*

**Figure S4.**
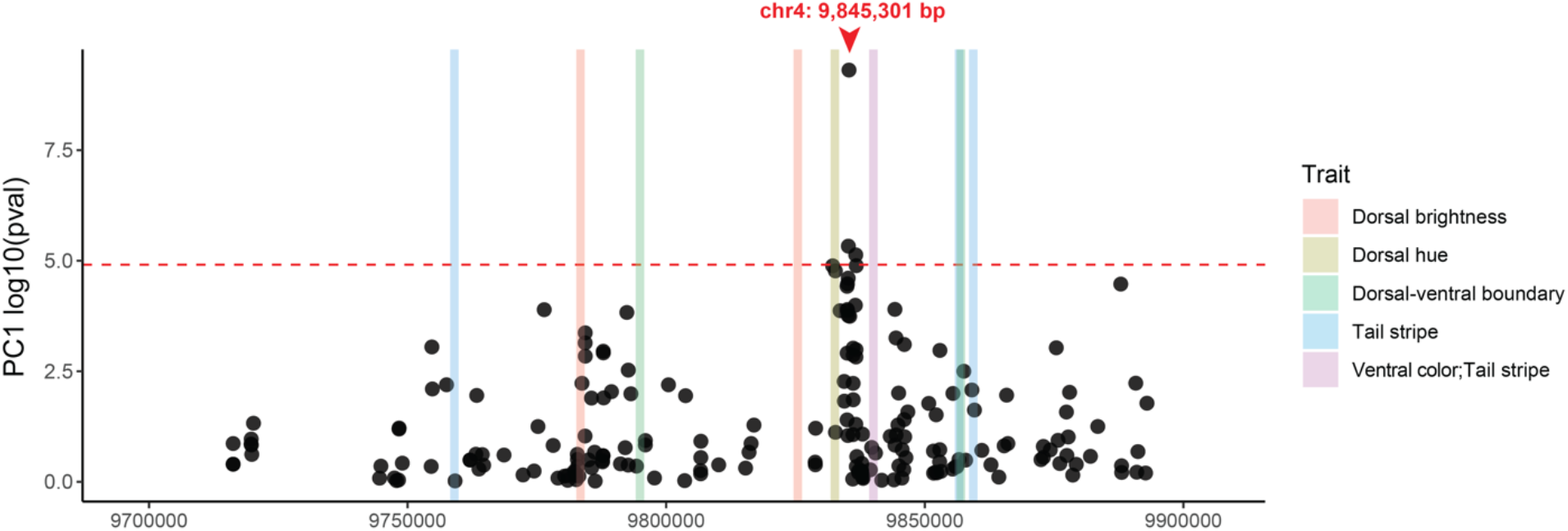
Location of the newly identified regulatory region in relation to previously implicated regions. The top-associated SNP (chr4:9,845,301) is shown in red. Vertical bars indicate regions of *Agouti* that are significantly associated with pigmentation traits in *P. maniculatus*, from Linnen et al. (2013).

**Figure S5.**
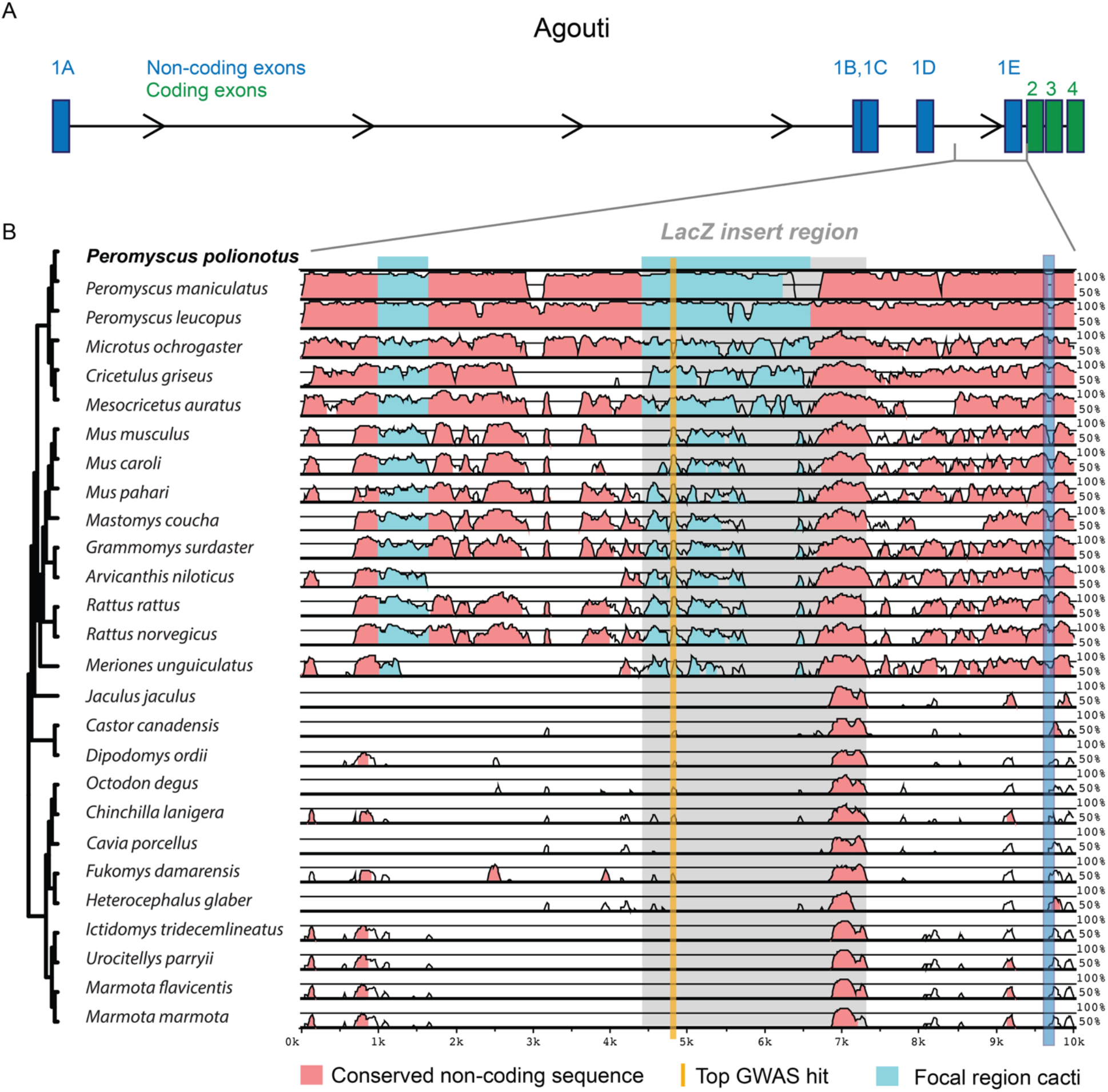
Sequence conservation in *Agouti* among 27 rodent species. 10 kb region encompassing the SNP with the strongest association to pPC1 (chr4:9,845,301) denoted in gold. Conserved regions are shown in pink. ‘Focal region cacti’ (light blue) indicate the regions identified by Saguaro (see Methods; Fig. 5) with a unique topology relative to the rest of the *Agouti* locus. The 2.6 kb region used in the *lacZ* reporter assay (grey) includes the cacti region (light blue), the top associated SNP (gold) and a conserved region (pink). One non-coding exon, 1E, is shown as a landmark (dark blue).

**Figure S6.**
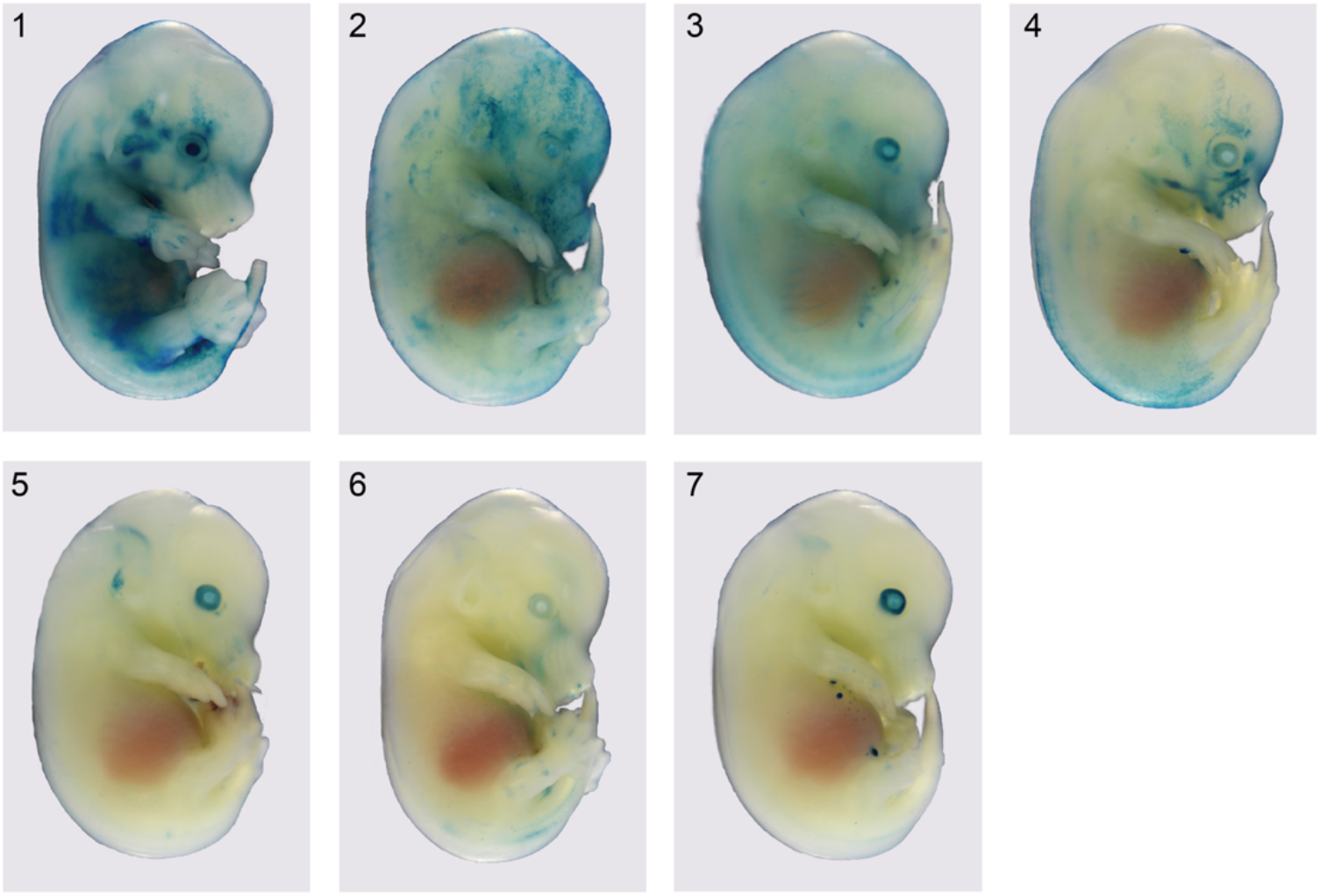
Transgenic embryos with visible skin *lacZ* expression produced by pronuclear injection of the candidate region-lacZ vector. All embryos are at stage E14.5, PCR-positive for the *lacZ* vector, and each represents an independent genomic integration event. Blue staining shows *lacZ* expression and regulatory element activity.

## Notes

### Competing Interest Statement

The authors have declared no competing interest.

